# Galectin-anchored indoleamine 2,3-dioxygenase suppresses local inflammation

**DOI:** 10.1101/2021.05.07.443161

**Authors:** Evelyn Bracho-Sanchez, Fernanda Rocha, Sean Bedingfield, Brittany D. Partain, Maigan A. Brusko, Juan M. Colazo, Margaret M. Fettis, Shaheen A. Farhadi, Eric Helm, Kevin Koenders, Alexander J. Kwiatkowski, Sabrina L. Macias, Antonietta Restuccia, Arun Wanchoo, Dorina Avram, Kyle D. Allen, Craig L. Duvall, Shannon M. Wallet, Gregory A. Hudalla, Benjamin G. Keselowsky

**Affiliations:** J. Crayton Pruitt Family Department of Biomedical Engineering, University of Florida, Gainesville, FL 32611; Department of Oral Biology, College of Dentistry, University of Florida, Gainesville, FL 32611; Department of Biomedical Engineering, Vanderbilt University, Nashville, TN, 37232; Department of Anatomy and Cell Biology, College of Medicine, University of Florida, Gainesville, FL 32610, USA; H. Lee Moffitt Cancer Center & Research Institute, Tampa, FL 33612; Division of Oral and Craniofacial Health Sciences, Adams School of Dentistry, Department of Microbiology and Immunology, School of Medicine, University of North Carolina at Chapel Hill, Chapel Hill, NC 27599

## Abstract

Chronic inflammation underlies the onset, progression and associated pain of numerous diseases.(*1*) Current anti-inflammatory treatments administered systemically are associated with moderate-to-severe side effects, while locally administered drugs have short-lived efficacy, and neither approach successfully modifies the underlying causality of disease.(*2*) We report a new way to locally modulate inflammation by fusing the enzyme indoleamine 2,3-dioxygenase 1 (IDO) to galectin-3 (Gal3). A general regulator of inflammation(*3*), IDO is immunosuppressive(*4*), catabolizing the essential amino acid tryptophan into kynurenine.(*5*) Recently we demonstrated that extracellular exogenous IDO regulates innate immune cell function(*6*), and envisioned delivering IDO into specific tissues would provide control of inflammation. However, proteins problematically diffuse away from local injection sites. Addressing this, we recently established that fusion to Gal3 anchors enzymes to tissues(*7*) via binding to extracellular glycans. Fusion protein IDO-Gal3 was retained in injected tissues and joints for up to a week or more, where it suppressed local inflammation in rodent models of endotoxin-induced inflammation, psoriasis, periodontal disease and osteoarthritis. Amelioration of local inflammation, disease progression and inflammatory pain were concomitant with homeostatic preservation of tissues without global immune suppression. Thus, IDO-Gal3 presents a new concept of anchoring immunomodulatory enzymes for robust control of focal inflammation in multiple disease settings.

---

Chronic inflammation, characterized by professional immune cell and resident tissue cell interactions, irreversibly damages tissues of the body and is an associated risk factor for a host of diseases such as cardiovascular disease, diabetes and cancer.(*1, 8*) A major challenge in the treatment of chronic inflammatory diseases is the development of therapeutics to safely and specifically direct resolution of inflammation in a site-specific manner.(*1*) Anti-inflammatory drugs such as glucocorticoids are pleiotropic, non-specifically affecting numerous pathways(*1*) and are consequently accompanied by issues of toxicity, resistance, and a wide array of serious adverse effects such as infection and defective wound healing.(*2*) Additionally, systemic immune modulation leads to disease states such as hypertension, osteoporosis, obesity, cataracts and diabetes.(*2*) Biologic immunosuppressive drugs functioning through either cytokine blockade, cell depletion or cell surface receptor blockade provide improved specificity, and can effectively modulate immune responses to halt disease progression in certain, but not all patients.(*9*) However, such treatments can also increase susceptibility to infections and disrupt tissue homeostasis, leading to cancer, exacerbation of congestive heart failure and neurologic events, among other pathologies.(*9*) Critically, each of these therapeutics require life-long continual use, and clinical options to resolve chronic inflammation and restore tissue homeostasis remain to be developed.(*10*)

The capability to direct cellular metabolism as a means to program immune responses has recently emerged as a profound new avenue for therapeutic immunomodulation.(*11*) Catabolism of the essential amino acid tryptophan (Trp) by the cytosolic enzyme indoleamine 2,3-dioxygenase 1 (IDO) and the resultant production of kynurenine metabolites is a general regulator of inflammation in response to sterile and pathogenic inflammatory stimuli.(*3, 5*) IDO catabolism of Trp is also a contributing factor to promoting fetal tolerance in pregnancy, warding off autoimmunity, and avoiding immune elimination in some forms of cancer.(*12*) Trp insufficiency via IDO activates the metabolic stress sensor general control nonderepressible 2 (GCN2) to regulate immune cell cycle(*13*) while kynurenine pathway metabolites activate anti-inflammatory programs, such as kynurenine binding to the aryl hydrocarbon receptor (AHR).(*14*) Additionally, the terminal product of the kynurenine pathway, nicotinamide adenine dinucleotide (NAD+), regulates innate immunity function in macrophages during aging and inflammation.(*15*) Informed by these data, we envisioned therapeutic delivery of exogenous IDO as a new and powerful regulator of chronic inflammatory diseases.

A major obstacle to treating localized tissue inflammation without systemic immunosuppression is the need to accumulate therapeutics at the intended site of action. Conceptually aligned with growth factors and antibodies engineered to bind extracellular matrix proteins for localized tissue retention(*16–19*), we recently demonstrated the utility of model enzymes fused to galectin-3 (Gal3), a carbohydrate-binding protein, as a generalizable means to restrict enzyme diffusion via binding to tissue glycans.(*7*) Gal3 binds N-acetyllactosamine and other β-galactoside glycans, as well as glycosaminoglycans, which are highly abundant in mammalian tissues(*20, 21*), collectively representing a more universal target than a specific extracellular matrix protein. Thus, Gal3 fusion to enzymes represents a promising approach to retain local enzymatic function at an intended site of action. Building upon this success, we engineered the IDO-Gal3 fusion with the expectation of creating a tissue-anchored IDO administered as a localized anti-inflammatory therapeutic (**Figure 1a,b**). Efficacy is described in four distinct inflammatory settings: endotoxin-induced inflammation, psoriasis, periodontal disease, and osteoarthritis.

**Figure 1.**
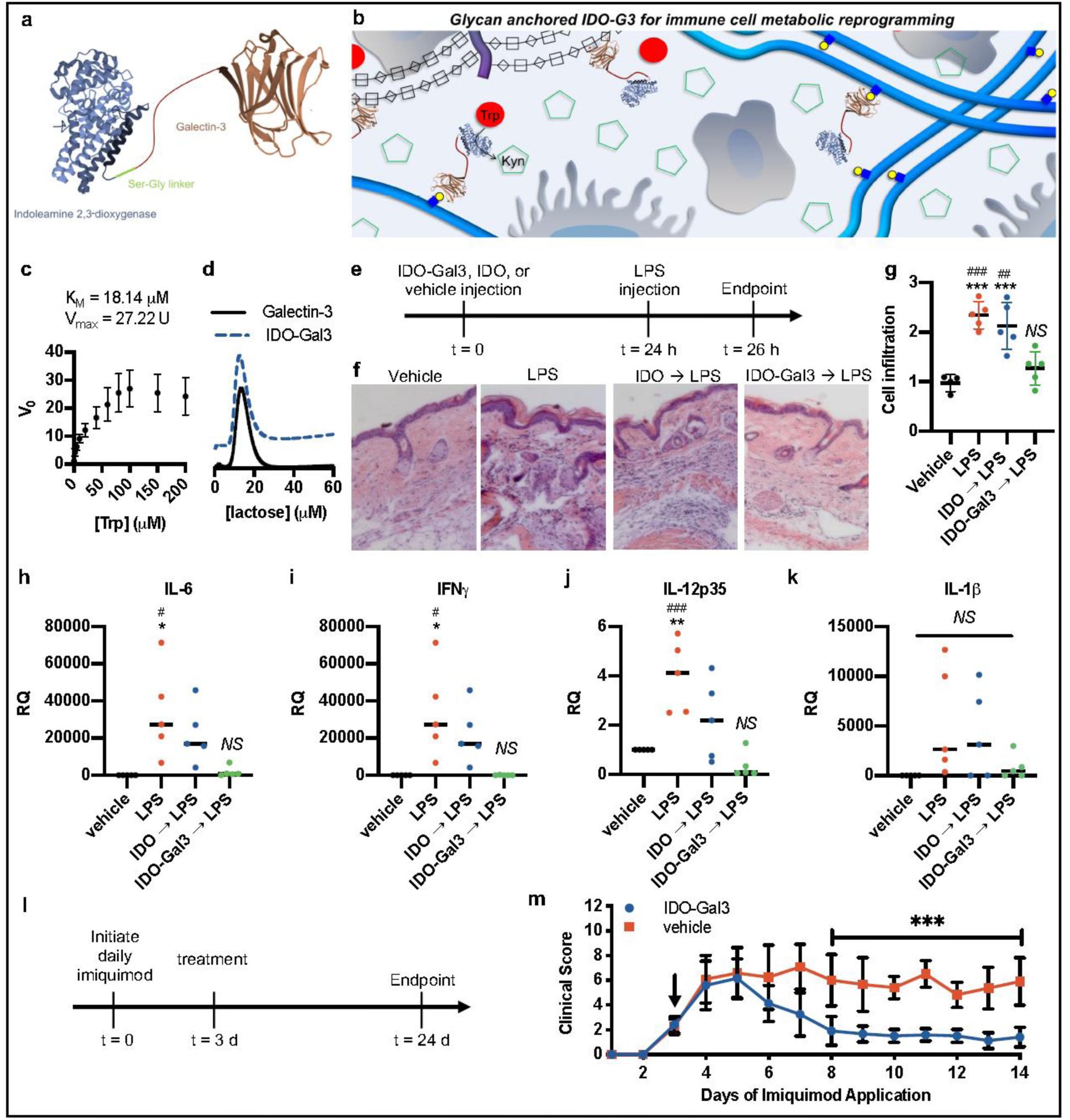
Indoleamine 2,3-dioxygenase anchored by galectin-3 (IDO-Gal3) suppresses inflammation. Schematic representation of (a) the IDO-Gal3 fusion protein and (b) the concept of anchoring IDO at different tissue locations through Gal3-mediated recognition of tissue glycans. Characterization of IDO-Gal3 (c) enzymatic activity and (d) binding to immobilized lactose. (e) Schedule to evaluate anchored IDO to suppress inflammation resulting from local lipopolysaccharide (LPS) injection. (f-g) Histological evaluation of IDO-Gal3 suppression of inflammation induced by injected LPS. (h-k) Quantification of inflammatory gene expression in tissues treated with anchored IDO prior to challenge with LPS. (l) Schedule and (m) clinical score to evaluate anchored IDO of inflammation induced by topical imiquimod. Statistical analysis: (g-k) One-way ANOVA with Tukey’s post-hoc. “NS” indicates no difference, two symbols denotes p < 0.01, three symbols denotes p < 0.001, n = 6; (m) Mann Whitney U-Test with alpha = 0.05, three symbols denotes p < 0.001, n = 12.

## IDO-Gal3 ameliorates skin inflammation

IDO-Gal3 catalyzed the conversion of Trp to kynurenine (**Figure 1c**), and bound immobilized lactose with comparable affinity as wild-type Gal3 (**Figure 1d**). Subcutaneous injection of IDO-Gal3 prevented inflammation induced by a local lipopolysaccharide (LPS) challenge better than IDO alone (**Figure 1e-k**). Histological analysis demonstrated that 24 h pretreatment with IDO-Gal3 mitigated the cellular infiltration that follows LPS challenge, whereas pretreatment with non-anchored IDO did not (**Figure 1e-g**). Naïve tissue cellular infiltration was unaffected by either IDO or IDO-Gal3 s.c. injection, compared to saline vehicle (PBS; **Supplementary Figure 1**). Pretreatment with IDO-Gal3 blocked transcription of the pro-inflammatory cytokines IL-6, IFN-γ, and IL-12p35, whereas expression was higher than vehicle control following pretreatment with IDO (**Figure 1h-j**). IL-1β transcript levels varied considerably following LPS challenge, but the trend suggested that pre-treatment with IDO-Gal3 also suppressed expression of this pro-inflammatory cytokine, whereas IDO did not (**Figure 1k**). Additionally, unchallenged naïve tissue gene expression of inflammatory cytokines was unaffected by either IDO or IDO-Gal3 s.c. injection, compared to saline vehicle (PBS; **Supplementary Figure 2**).

Local injection of IDO-Gal3 also resolved inflammation in a murine psoriasis model (**Figure 1l-m**). Topical daily application of imiquimod over 14 d induced inflammation which peaked at day 5, reflected by psoriasis severity clinical score. Vehicle injection did not affect the severity score, which remained elevated for the remaining 14 d period. In contrast, subcutaneous injection of IDO-Gal3 on day 3 led to a dramatic decrease in clinical score by day 8, which remained low for the duration, despite continued daily imiquimod challenge. In sum, IDO-Gal3 ameliorated skin inflammation both prophylactically and therapeutically, with different inflammatory insults acting through different specific receptors (CD14/TLR4 for LPS, and TLR7 for imiquimod).(*22*)

## IDO-Gal3 functions locally and is not systemically suppressive

We recently demonstrated Gal3 fusion to various enzymes endows glycan-binding affinity that restricts enzyme diffusion in tissues, prolonging retention and localizing enzymatic catalysis.(*7*) In vivo imaging detected fluorophore-labeled IDO-Gal3 out to 14 d with s.c. hock injection site (**Supplementary Figure 3a**). Increased kynurenine levels in surrounding tissue detected via mass spectrometry analysis were consistent with in vivo IDO enzymatic function (**Supplementary Figure 3b**). Further, bioluminescence imaging of anchored reporter enzyme, the fusion of NanoLuc^TM^ luciferase and Gal3 (NL-Gal3) was retained at high levels compared non-anchored NanoLuc^TM^, and suggested NL-Gal3 injected into s.c. hock was largely restricted to the injection site, although lower enzyme activity was also detected in the adjacent tibial region, in plasma and the excretory pathways of the urine and feces (**Supplementary Figure 4**). To determine if localized IDO-Gal3 induced systemic immunosuppression, 120 h after injection, mice were subjected to LPS challenge at ipsilateral and contralateral sites, as well as an oral *Listeria monocytogenes* infection challenge (**Supplementary Figure 3c-k**). Prophylactic administration of IDO-Gal3 in the ipsilateral hock blocked LPS-induced inflammatory cell infiltration 120 h after pretreatment, whereas infiltrate was elevated at the contralateral LPS challenge site, suggesting the inflammatory blockade was localized to the ipsilateral site (**Supplementary Figure 3c-d**). Administering IDO-Gal3 into the s.c. hock did not alter *L. monocytogenes* clearance in the liver or spleen when compared to vehicle control (**Supplementary Figure 3e-g**), which indicated maintenance of a globally functioning immune system concurrent with the tissue-localized suppression. In corroboration with cellular infiltration data, even 120 h after pretreatment with IDO-Gal3 (**Supplementary Figure 3c-d**), expression of IL-6 and IL-1β at the ipsilateral LPS challenge site (**Supplementary Figure 3h-i**) was suppressed, as was expression of IL-12p35, IL-12p40 and IFN-γ (**Supplementary Figure 5**), which prevented increased plasma IL-6 protein (**Supplementary Figure 3j**) and maintained baseline plasma IL-1β levels (**Supplementary Figure 3k**) detected in serum by mass spectrometry. Collectively, these observations suggested tissue-anchored IDO-Gal3 induced potent local suppression of inflammation, without globally suppressing immune function.

## IDO-Gal3 prevents and inhibits periodontal disease progression

Periodontal disease is a class of non-resolving chronic inflammatory diseases resulting from localized mucosal inflammation and osteoclast activation, and leading to soft (mucosa) and hard (bone) tissue destruction.(*23*) Submandibular injection of IDO-Gal3 was locally retained, suppressed mucosal inflammation, and spared alveolar bone loss (**Figure 2**), evaluated with both a prophylactic and therapeutic administration scheme in a polymicrobial periodontal disease mouse model(*24*)(**Figure 2a-b**). In vivo imaging of anchored reporter NL-Gal3 demonstrated gingival retention for 120 h (**Figure 2c-d**) whereas tissue localization for the non-anchored NL could not be measured after 36 h (**Figure 2d**). Retention time was not affected by the presence or absence of polymicrobial infection (**Figure 2e**). The primary outcome of periodontal disease, mandibular bone loss, was quantified by micro-CT analysis. When administered prophylactically (IDO-Gal3-P) submandibular injection of IDO-Gal3 prevented bone loss and inhibited further bone loss when administered therapeutically (IDO-Gal3-T; **Figure 2f**). Specifically, loss of trabecular bone volume (**Figure 2g**) and thickness (**Supplementary Figure 6**), along with vertical bone loss (**Figure 2h**) were prevented by either prophylactic or therapeutic administration of IDO-Gal3. IDO-Gal3 also inhibited gingival inflammation as measured by protein levels of cytokines IL-6 (**Figure 2i**) and IL-1β (**Figure 2j**), and chemokine MCP-1 (**Figure 2k**). In contrast, IL-10 production was largely maintained through IDO-Gal3 treatment (**Figure 2l**), whereby levels were closer to uninfected mice, suggesting a return to homeostatic control. A more pronounced diminution of bone loss and suppression of inflammation was observed upon therapeutic administration of IDO-Gal3 when compared to prophylactic administration (**Figure 2h-k**), where the inflammatory cytokines IL-12p70, IP-10, KC-17, MIP-2, and IL-33 were most affected (**Supplementary Figure 7**). IDO-Gal3 administered to uninfected mice (IDO-Gal3-U) had negligible effect on mandibular bone and production of inflammatory cytokines, however, increased levels of IL-10, and to a smaller extent, MCP-1, were observed (**Figure 2j-l**). Furthermore, IDO-Gal3 administration did not affect the amount of either *P. gingivalis* or *A. actinomycetemcomitans* recovered from the gingiva, regardless of prophylactic or therapeutic administration scheme (**Supplementary Figure 8**). Together these data indicate that IDO-Gal3 addresses the clinical therapeutic requirements for treatment of periodontal disease: localized regulation of inflammation without enhancing bacterial accumulation, prevention of continued mandibular bone loss even after disease progression has begun.

**Figure 2.**
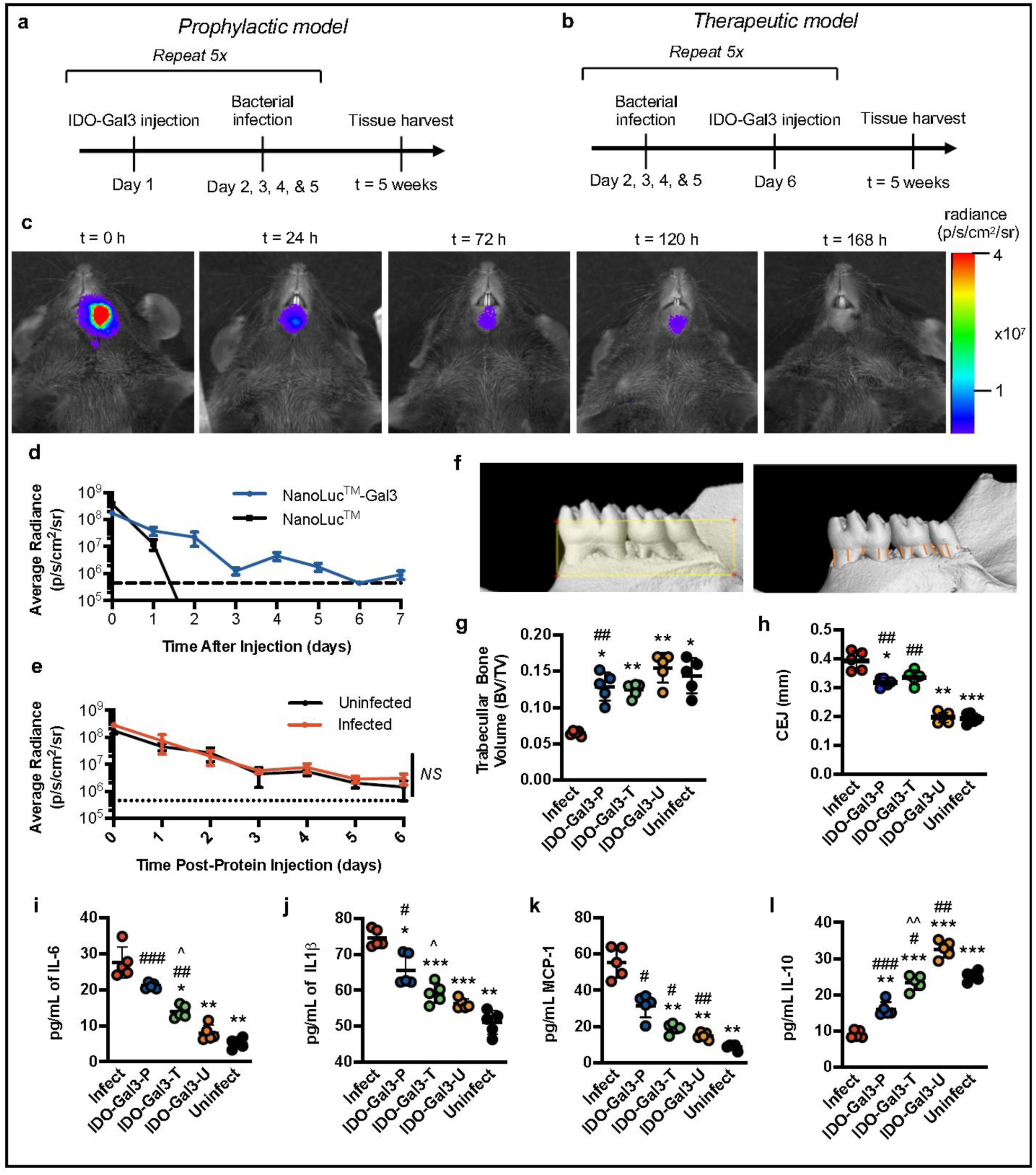
IDO-Gal3 prevents and inhibits periodontal disease progression. Efficacy was investigated using a polymicrobial mouse model of periodontal disease, and either (a) prophylactic or (b) therapeutic administration. (c-d) Anchored-fusion reporter enzyme NanoLuc^TM^-Gal3 (NL-Gal3) injected submandibular was retained locally for 120 h, by in vivo imaging, in contrast to non-anchored NanoLuc^TM^ which diffused away from the injection site (d). (e) Infection state did not alter submandibular retention time of NL-Gal3. (f) Representative images of micro-CT analysis quantifying (g) trabecular bone volume and (h) horizontal bone loss, by measuring the cemento-enamel junction (CEJ). Submandibular injection of IDO-Gal3 blocked mandibular bone loss in prophylactic (IDO-Gal3-P) administration, and halted mandibular bone loss in therapeutic (IDO-Gal3-T) administration. IDO-Gal3 reduced gingival inflammatory protein levels of IL-6 (i) and IL-1β (j) and MCP-1 (l), with a more pronounced effect from therapeutic administration (IDO-Gal3-T). (k) IDO-Gal3 induced the expression of IL-10 protein, restoring levels closer to that of the uninfected control. Statistical analysis: (e) Student’s t-test, ns = no difference, n = 5; (g-l) * sig diff compared to infect, # = significant difference compared to Uninfect, ^ = difference between Prophylactic and Therapeutic, ANOVA w/ Tukey’s. One symbol denotes p < 0.05, two symbols denotes p < 0.01, three symbols denotes p < 0.001, n = 5.

## IDO-Gal3 protects against developing post-traumatic osteoarthritis

Post-traumatic inflammatory osteoarthritis (PTOA), a form of osteoarthritis (OA), commonly arises after a ligament or meniscus tear, or following repeated over-loading of a joint. A cyclic mechanical overloading PTOA mouse model(*25, 26*) was utilized to test for ability of IDO-Gal3 to reduce load-induced inflammatory joint damage. Intra-articular injection of IDO-Gal3 was locally retained, suppressed inflammation, and spared joint tissue destruction (**Figure 3**). Cyclic mechanical loading was applied to knees of aged mice (**Figure 3a; Supplementary Figure 9**), with mechanical damage and consequent inflammation resulting in synovial inflammation and cartilage degradation. In a retention study, fluorescently-labeled IDO or labeled IDO-Gal3 was administered after two weeks of initial joint mechanical loading. Imaging at 24 h (**Supplementary Figure 10**) and day 7 post-injection revealed that retention of labeled IDO-Gal3 in the joint was significantly higher than non-anchored IDO, visualized in explanted knees (**Figure 3b**), and quantified (**Figure 3c**) by ex vivo image analysis. The pharmacokinetic area-under-curve, calculated from daily intravital imaging over 7d, was approximately two-fold higher for IDO-Gal3 than IDO alone (**Figure 3d**). Weekly intra-articular administration of IDO-Gal3 blocked gene expression of inflammatory cytokines IL-6 and IL-12p40 in both articular cartilage (**Figure 3e-f**) and draining lymph nodes (**Figure 3g-h**), whereas IDO alone did not. In contrast, expression of neither MMP13 nor TNF-α was modulated, at either location (**Supplementary Figure 11**). Critically, IDO-Gal3 reduced load-induced histological changes, as scored by a treatment-blinded histopathologist by both total joint health score (degenerative joint disease (DJD) score) (**Figure 3i-j**) and articular cartilage score (OARSI) (**Figure 3k-l**). Additional histological specimens support these trends (**Supplementary Figure 12-15**), and scoring criteria description is provided in **Supplementary Figure 16-17**. In sum, Gal3 fusion provided local joint retention, and IDO-Gal3 potently reduced inflammatory gene expression and mechanical overloading-induced structural changes in the joint.

**Figure 3.**
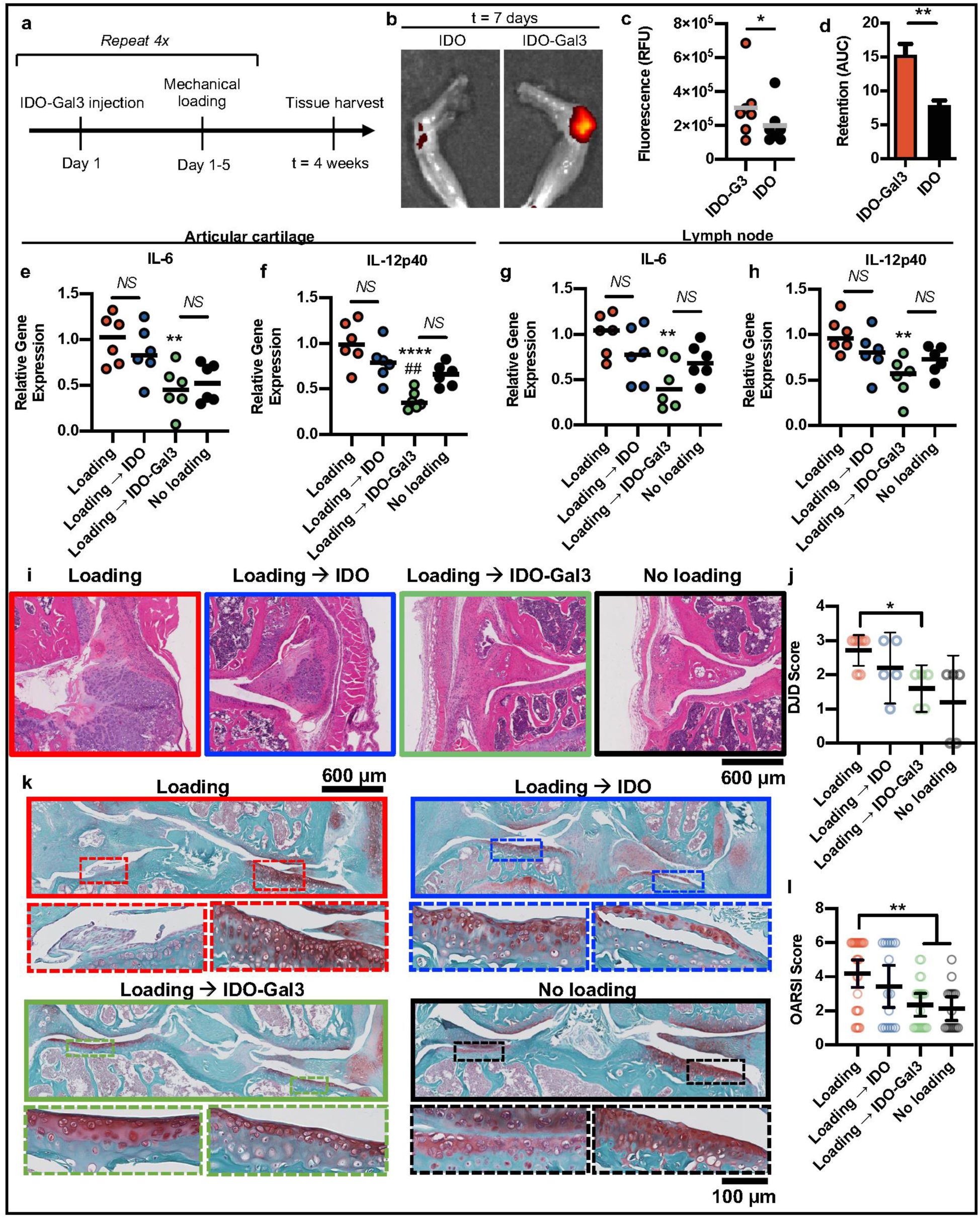
IDO-Gal3 protects against upregulation of inflammatory gene expression and joint structural changes from load-induced osteoarthritis. A 4 week cyclic mechanical overloading PTOA mouse model was utilized in mice treated weekly with intra-articular IDO-Gal3 injections (a). Knee explant fluorescence imaging illustrating (b) and quantifying (c) day 7 retention of fluorescently-labeled IDO and IDO-Gal3, administered after 2 weeks of mechanical loading. Area-under-curve pharmacokinetic analysis from daily fluorescent imaging over 7 d (d). Gene expression of cytokines IL-6 and IL-12p40 was measured by quantitative PCR in articular cartilage (e-f) and draining lymph nodes (g-h). H&E stained joint histology on knees highlighting synovial inflammation and thickening (i) and degenerative joint disease (DJD) scoring of joint disease progression (j). Safranin-O histological stain highlighting structure changes and proteoglycan content of the articular cartilage (k) and OARSI scoring of cartilage structure (l). Statistical analysis: (c, d) student’s t-test, * denotes p < 0.05, ** denotes p < 0.01, n = 6; (e-h) ANOVA with Tukey’s post-hoc, “NS” indicates no difference between groups, ** denotes p < 0.01, n = 6; (j, l) Brown-Forsythe and Welch ANOVA, “NS” indicates no difference between groups, * denotes p < 0.05, ** denotes p < 0.01, n = 6.

## IDO-Gal3 reduces inflammation and pain, improving gait in established PTOA osteoarthritis

Osteoarthritis pain and disability are associated with chronic inflammation in an irreparably damaged joint.(*27*) In order to investigate a single-dose IDO-Gal3 treatment in established OA, a surgically-induced PTOA rat model was utilized, where OA developed over 8 weeks of accrued joint damage following transection of the medial collateral ligament and medial meniscus (MCLT+MMT)(*28*). Intra-articular injection of IDO-Gal3 was locally retained, suppressed joint inflammation, reduced OA-associated pain and improved rat gait (**Figure 4**). Administered into both OA and contralateral healthy joints (**Figure 4a**), luminescence of the NL-Gal3 anchored reporter was measurable over multiple days post-injection while non-anchored NL was not found in the joint after 1 d (**Figure 4c**). Joint residence times (95% decay) for NL-Gal3 were 2 d in OA joints and 1.3 d in healthy joints, which is up to 4-fold longer, compared to 0.5 d for NL in both OA and healthy joints (**Figure 4c**, **Table**). Injection of IDO-Gal3 at 8 w post-surgery (**Figure 4b**) blocked local inflammatory cytokine production, and furthermore reversed OA-associated tactile sensitivity and compensatory gait (**Figure 4d-g**). Treatment with IDO-Gal3 reduced protein production of inflammatory cytokine IL-6 in OA joints to levels found in healthy joints (**Figure 4d**). Additionally, local levels of inflammatory chemokine MCP-1 followed this same trend (p = 0.09; **Figure 4e**). Tactile sensitivity as a metric of OA-associated pain was quantified over 3 wk using von Frey monofilaments (**Supplementary Figure 18**, absolute values). Plotted as the change in paw withdrawal threshold (post-injection – pre-injection), treatment with IDO-Gal3 reversed limb hypersensitivity by post-injection day 8, persisting through day 23, while saline vehicle injection provided only marginal improvement (**Figure 4f**). Rodent gait was evaluated by simultaneous high-speed videography and force plate recordings(*29*). IDO-Gal3-treated rats utilized faster walking speeds on post-injection day 16, and tended to walk faster on post-injection day 23 (**Supplementary Figure 19a-b**). Notably, peak vertical force plotted as a function of velocity demonstrated that a single IDO-Gal3 injection corrected imbalanced weight distribution in OA-affected animals, where dynamic loading on IDO-Gal3 treated limbs was indistinguishable from healthy contralateral controls throughout the study duration (**Figure 4g**, **top row**; **Supplementary Figure 19c**). This contrasted with the distinct loading imbalance between saline-treated and contralateral limbs (**Figure 4g**, **bottom row**; **Supplementary Figure 19c**). Notably, IDO-Gal3 and saline treatments were significantly different at post-injection days 9, 16 and 24. Altogether, these data indicate potent amelioration of established OA inflammation, along with concomitant improvement in OA symptoms.

**Figure 4.**
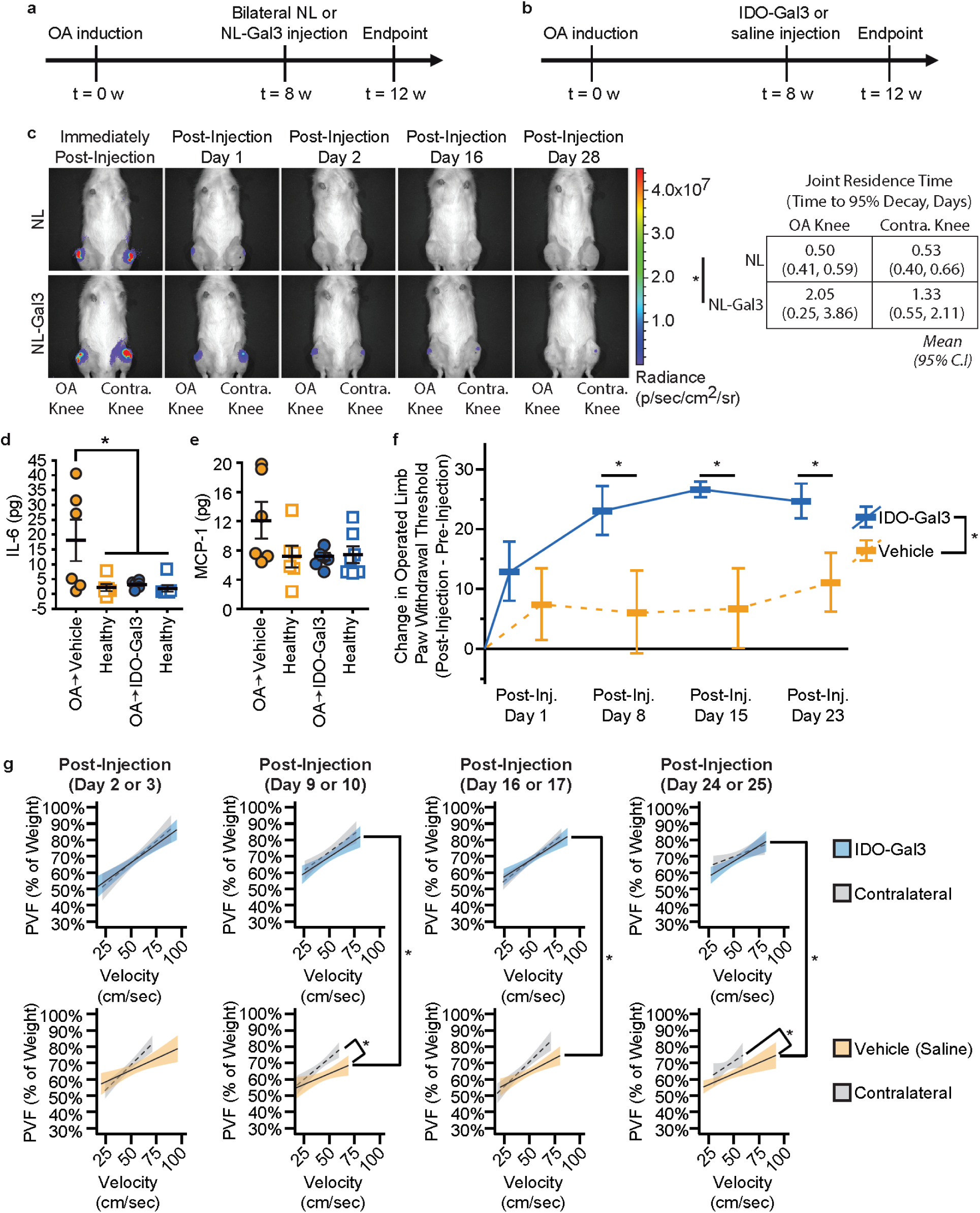
IDO-Gal3 reduces inflammation, decreases tactile hypersensitivity and improves gait after established osteoarthritis. Following a surgically-simulated meniscal tear in the rat, clearance of NL-Gal3 was assessed 8 weeks after accrued joint damage (a). Using the same model and experimental design, the effects of IDO-Gal3 on OA-associated inflammation and symptoms were evaluated (b). Luminescence imaging demonstrated NL-Gal3 is retained in both OA-affected and healthy joints longer than unconjugated NL, with significantly longer times to 95% decay in NL-Gal3 injected knees (c). At 28 days after injections in OA-affect joints, saline-treated knees had elevated levels of IL-6 (p≤0.03), while IDO-Gal3 levels were comparable to healthy contralateral knees (d). Intra-articular levels of MCP1 (i.e., CCL2) followed a similar profile, but CCL2 was not significantly elevated in saline treated knees for this study (p=0.09, e). Relative to saline-treated animals, IDO-Gal3-treated animals had improvements in tactile sensitivity levels across time (p=0.03, f). Raw data are shown in Supplementary Figure 18-19. IDO-Gal3 treated animals had similar weight distribution between their OA-affected and contralateral limb, while saline treated animals had significant off-loading of the OA-affected limb (g). Behavioral data is a linear mixed affects model treating repeated measures within an animal across time as a random effect. Everything else is a 1-way ANOVA with paired samples. All post-hoc tests are comparisons of estimated means using a Tukey’s HSD adjustment for compounding type 1 errors, * denotes p < 0.05.

In sum, the IDO-Gal3 tissue-anchored enzyme fusion represents a potent new class of anti-inflammatory protein therapeutic with broad potential implications for better treatment outcomes and reduced systemic side effects for a broad variety of localized, inflammatory diseases. The cytosolic immunomodulatory enzyme IDO was newly conceived as an exogenously functioning and locally deliverable protein therapeutic to directly manipulate metabolism of the essential amino acid tryptophan, acting here as a prodrug, systemically available. Fusion with Gal3 prolonged enzyme localization and retention through binding to glycans, which are ubiquitously expressed across mammalian tissues. The conservation of glycans across species enables translation without requiring redesigning for human-specific targets. Administered IDO-Gal3 robustly controlled local inflammation across four distinct inflammatory settings in multiple species, and in multiple tissue environments. Treatment restored homeostasis, preserved tissue integrity and function, and relieved inflammation-associated pain. Administration of IDO-Gal3 uniformly blocked IL-6 in each inflammatory model investigated, among numerous other cytokines and chemokines, and did not induce systemic immunosuppression. Altogether, this work highlights how the galectin-anchored enzyme overcomes the drug delivery obstacle of diffusion of a therapeutic away from intended site of action, and points to IDO-Gal3 as a powerful new anti-inflammatory drug.

**Supplementary Information** is available for this paper.

**Correspondence** and requests for materials should be addressed to BGK and GAH.

## Acknowledgment

This work was supported by the National Institutes of Health (R01 DE027301 to BGK, SMW and GAH; R01 DK091658 and R01 DK098589 to BGK; R01 AI133623 to BGK and DA; R03 EB019684 and R35 GM133697 to GAH; R01 AR068424 and R01 AR071431 to KDA; T32 DK108736 to AJK; UL1TR001427 and TL1TR001428 to SAF), by the Department of Defense (CDMRP OR130302 to CLD), and by the National Science Foundation (DMR-1455201 to GAH).

## Author contributions

EB-S, FR, BDP, SB, SLM, and AJK contributed to the design of the work, the acquisition, analysis, and interpretation of data. AR, MMF, SAF, EH, JMC, KK, MAB, AW contributed to the acquisition, and analysis of data. DA contributed to the design of the work, and the analysis and interpretation of data. CLD, KDA, SMW contributed to the design of the work, and the analysis and interpretation of data, and substantively revised the manuscript. BGK and GAH conceived of and directed the work, contributed to the design of the work, contributed to analysis and interpretation of data, and wrote the manuscript. All authors contributed to and approved the submitted version.

## Methods

### Recombinant protein expression and purification

NanoLuc™ is the tradename of an engineered deep sea shrimp luciferase variant developed by Promega Corporation.(*30*) Genes encoding fusion proteins were inserted into pET-21d(+) vectors between NcoI and XhoI sites. IDO-Gal3 genetic and protein sequences are provided in Supplementary Information. Plasmids confer ampicillin resistance and were first transformed into One Shot™ TOP10 Chemically Competent *E. coli* (ThermoFisher) and selected on Luria-Bertani (LB) ampicillin (50 μg/mL) agar, incubated overnight at 37 °C. Isolated colonies from the plates were subcultured in 5mL LB ampicillin broth overnight at 37 °C with orbital shaking. Successful transformants were confirmed by Sanger sequencing (Genewiz). Positive DNA sequences were then transformed into the kanamycin-B resistant expression strain, Origami™ B (DE3) *E. coli* (Novagen) and selected on LB ampicillin with kanamycin B (15 μg/mL) agar. Positive clones were picked, banked in LB with 10% w/v glycerol, and used to preculture 5 mL of LB ampicillin and kanamycin. Overnight precultures were used to inoculate 1 L 2×TY media (16 g tryptone, 10 g yeast extract, 5 g NaCl) with ampicillin and kanamycin B at 37 °C, 225 rpm in an orbital shaker until approximate exponential growth phase (O.D.600 nm = 0.6–0.8). IDO-Gal3 expression cultures were supplemented with 500 μM δ-Aminolevulinic acid (Sigma) at the time of inoculation. Recombinant protein expression was triggered using 0.5mM isopropyl β-D-1-thiogalactopyranoside (ThermoFisher) and incubated for 18 h in an orbital shaker at 18 °C. Bacteria were washed and pelleted with PBS via centrifugation (13,180 × g at 4 °C for 10 min) with a superspeed centrifuge (ThermoFisher). Afterwards, the pellet was weighed, resuspended in 4 mL PBS/gram pellet with protease inhibitor (ThermoFisher), and was disrupted by sonic dismembration (15 seconds on, 45 seconds off, 10 minute cycling time, ThermoFisher). Following dismembration, lystates were treated with 20 units/gram pellet DNAse I (ThermoFisher), and 800 μg/gram pellet lysozyme (ThermoFisher). The lysate was centrifuged (38,360 × g at 4 °C for 15 min) to remove the insoluble fraction and the supernatant was collected by decanting. Supernatant containing soluble recombinant protein was loaded into α-lactose agarose (Sigma) affinity column preequilibrated with PBS. Columns were washed with 20 column volumes PBS and recombinant proteins were eluted with 100 mM β-d-lactose (Sigma) prepared in PBS. A final polishing step was performed by 200 kDa size exclusion chromatography on AKTA pure chromatography system (GE Life Sciences) to remove α-lactose and further isolate IDO-Gal3. Protein purity was determined by sodium dodecyl sulfate polyacrylamide gel electrophoresis (SDS-PAGE) and Coomassie staining. Endotoxin contaminants were removed by endotoxin removal solution (Sigma) following manufacturer’s instructions. Endotoxin content was analyzed using Chromo-LAL kinetic chromogenic endotoxin quantification assay (Associates of Cape Cod Inc.), and determined to be below 0.1 EU/mL in all stocks.

### IDO Enzymatic Activity Assay

Recombinant human IDO expressed in Escherichia coli was purchased from R&D Systems (Minneapolis, MN) with a predicted molecular mass of 46 kDa a specific activity of >500pmoles/min/μg (or for equimolar calculations >29 pmol NFK/min/pmol IDO) as measured by its ability to oxidize L-tryptophan to N-formyl-kynurenine (NFK). The specific activity of both proteins, IDO and IDO-Gal3, was measured before experiments to ensure maximal effect at the beginning of the assay following the IDO manufacturer’s protocol, IDO-Gal3 was reacted in equimolar amounts to IDO in the standard protocol, and activities were compared using the unit pmol NFK/min/pmol IDO to more accurately compare activity.

### IDO-Gal3 Binding Affinity

Affinity of IDO-Gal3 for lactose was determined using affinity chromatography in an AKTA Pure chromatography system (GE Life Sciences) equipped with consumer-packable glass column (GE Life Sciences) packed with α-lactose agarose affinity resin (Sigma-Aldrich). Proteins were eluted with a linear gradient of β-lactose (Sigma-Aldrich) in phosphate buffer.

### IDO-Gal3 DNA Sequence: (Restriction sites Underlined)

<u>CCATGG</u>CGCACGCGATGGAAAACAGCTGGACCATCAGCAAAGAGTACCACATTGACGAGGA AGTTGGTTTCGCGCTGCCGAACCCGCAGGAAAACCTGCCGGACTTCTATAACGATTGGATGTTTATCGCGAAGCACCTGCCGGATCTGATTGAGAGCGGCCAGCTGCGTGAGCGTGTGGAAAAACTGAACATGCTGAGCATCGACCACCTGACCGATCACAAGAGCCAACGTCTGGCGCGTCTGGTTCTGGGTTGCATTACGATGGCGTACGTGTGGGGCAAAGGTCACGGCGACGTGCGTAAGGTTCTGCCGCGTAACATCGCGGTTCCGTACTGCCAACTGAGCAAGAAACTGGAACTGCCGCCGATTCTGGTGTATGCGGACTGCGTTCTGGCGAACTGGAAGAAGAAGGACCCGAACAAACCGCTGACCTATGAGAACATGGATGTGCTGTTCAGCTTTCGTGACGGTGATTGCAGCAAGGGCTTCTTTCTGGTGAGCCTGCTGGTTGAAATCGCGGCGGCGAGCGCGATCAAAGTGATTCCGACCGTTTTCAAGGCGATGCAGATGCAAGAGCGTGACACCCTGCTGAAAGCGCTGCTGGAAATCGCGAGCTGCCTGGAGAAGGCGCTGCAGGTGTTTCACCAAATTCACGATCACGTTAACCCGAAAGCGTTCTTTAGCGTGCTGCGTATCTACCTGAGCGGTTGGAAGGGCAACCCGCAGCTGAGCGACGGTCTGGTTTATGAGGGCTTCTGGGAAGATCCGAAAGAGTTTGCGGGTGGCAGCGCGGGTCAGAGCAGCGTGTTCCAATGCTTTGACGTTCTGCTGGGCATTCAGCAAACCGCGGGTGGCGGTCATGCGGCGCAGTTCCTGCAAGATATGCGTCGTTACATGCCGCCAGCGCACCGTAACTTCCTGTGCAGCCTGGAAAGCAACCCGAGCGTGCGTGAGTTTGTTCTGAGCAAAGGTGACGCGGGCCTGCGTGAAGCGTATGATGCGTGCGTGAAGGCGCTGGTTAGCCTGCGTAGCTACCACCTGCAGATCGTTACCAAATATATCCTGATTCCGGCGAGCCAGCAACCGAAAGAAAACAAGACCAGCGAGGACCCGAGCAAACTGGAGGCGAAGGGTACCGGCGGTACCGATCTGATGAACTTTCTGAAG ACCGTGCGTAGCACCACCGAGAAGAGCCTGCTGAAAGAGGGT<u>GGATCC</u>GGCGGCGGCAGCG GCGGCAGCGGCGGCAGCGGCGGC<u>GAATTC</u>GCGGACAACTTCAGCCTGCACGATGCGCTGAG CGGTAGCGGTAACCCGAACCCGCAGGGTTGGCCGGGTGCGTGGGGTAACCAACCGGCGGGTGCGGGTGGCTACCCGGGTGCGAGCTATCCGGGTGCGTATCCGGGTCAGGCTCCGCCGGGTGCGTACCCGGGCCAAGCTCCGCCGGGTGCTTATCCTGGTGCGCCGGGCGCGTACCCGGGTGCGCCGGCGCCGGGCGTGTACCCGGGTCCGCCGAGCGGTCCGGGCGCGTATCCGAGCAGCGGCCAGCCGAGCGCGCCGGGTGCGTATCCGGCGACCGGCCCGTATGGTGCGCCGGCGGGTCCGCTGATTGTTCCGTATAACCTGCCGCTGCCGGGTGGCGTGGTTCCGCGTATGCTGATCACCATTCTGGGCACCGTGAAGCCGAACGCGAACCGTATCGCGCTGGACTTCCAACGTGGTAACGATGTTGCGTTCCACTTTAACCCGCGTTTTAACGAGAACAACCGTCGTGTGATTGTTTGCAACACCAAACTGGACAACAACTGGGGCCGTGAGGAACGTCAGAGCGTGTTCCCGTTTGAGAGCGGCAAGCCGTTCAAAATTCAAGTGCTGGTTGAACCGGACCACTTTAAGGTGGCGGTTAACGATGCGCACCTGCTGCAGTACAACCACCGTGTTAAGAAACTGAACGAAATCAGCAAACTGGGCATCAGCG GTGACATTGATCTGACCAGCGCGAGCTATAACATGATT<u>CTCGAG</u>

### Amino Acid Sequence

MAHAMENSWTISKEYHIDEEVGFALPNPQENLPDFYNDWMFIAKHLPDLIESGQLRERVEKLNMLSIDHLTDHKSQRLARLVLGCITMAYVWGKGHGDVRKVLPRNIAVPYCQLSKKLELPPILVYADCVLANWKKKDPNKPLTYENMDVLFSFRDGDCSKGFFLVSLLVEIAAASAIKVIPTVFKAMQMQERDTLLKALLEIASCLEKALQVFHQIHDHVNPKAFFSVLRIYLSGWKGNPQLSDGLVYEGFWEDPKEFAGGSAGQSSVFQCFDVLLGIQQTAGGGHAAQFLQDMRRYMPPAHRNFLCSLESNPSVREFVLSKGDAGLREAYDACVKALVSLRSYHLQIVTKYILIPASQQPKENKTSEDPSKLEAKGTGGTDLM NFLKTVRSTTEKSLLKEG<u>GS</u>GGGSGGSGGSGG<u>EF</u>ADNFSLHDALSGSGNPNPQGWPGAWGNQP AGAGGYPGASYPGAYPGQAPPGAYPGQAPPGAYPGAPGAYPGAPAPGVYPGPPSGPGAYPSSGQPSAPGAYPATGPYGAPAGPLIVPYNLPLPGGVVPRMLITILGTVKPNANRIALDFQRGNDVAFHFNPRFNENNRRVIVCNTKLDNNWGREERQSVFPFESGKPFKIQVLVEPDHFKVAVNDAHLLQYNHRVKKLNEISKLGISGDIDLTSASYNMILEHHHHHH

### LPS Administration at the Hock as a Model of Inflammation

B6 mice were administered 2.1 μg of IDO-Gal3 in 40 μL of PBS subcutaneously (ipsilateral and contralateral) at the region of the hock and challenged with 2 ng/g of lipopolysaccharide (LPS) in 40 μL, after 24 h or 120 h post IDO-Gal3 treatment, to assess local and distal modulation of inflammation. 2 h after LPS administration animals were euthanized and the injection site collected.

### Quantitative PCR

Soft tissue was separated from bone and stored in RNAlater RNA Stabilization Reagent (Qiagen) in preparation for qPCR. Soft tissues were homogenized and RNA purified using RNeasy Protect Mini Kit (Qiagen). cDNA was synthesized from RNA using the High-Capacity cDNA Reverse Transcriptase Kit (ThermoFisher) for use in qPCR in accordance with manufacturers instructions. qPCR analysis was ran with primers specific for pro-inflammatory cytokines (I12a, Il12b, Il1b, Ifng and Il6). Results are presented as the ratio of gene expression to GAPDH expression determined by the relative quantification method. Treatment groups were normalized to PBS only group.

### Hock Infiltration

Tissues were fixed with 10% Neutral Formalin pH 7.4 overnight. After fixation, tissues were washed in diH2O and decalcified by storing in 10% Ethylenediaminetetraacetic acid (EDTA) at 4°C for 3 weeks. Samples were assessed every 2-3 days for stiffness and EDTA solution replenished. Tissues were submitted in 70% ethanol to the University of Florida Molecular Pathology Core for processing, paraffin embedding, sectioning, mounting and staining with Haemotoxylin and Eosin (H&E). Tissues were imaged using a Zeiss Axiovert 200M with a 20X objective lens through the multidimensional acquisition module. 11 images were taken per tissue and scored by two blinded independent individuals based on cellular infiltration (0: absent, 1:mild, 2:moderate, 3:severe) and epidermis hypertrophy (0:0-20 μm in thickness, 1: 21-40 μm, 2: 41-60 μm, 3: 61μm or above).

### Mass Spectrometry

2.2μg of IDO-Gal3 in 40 μL of PBS or 40 μL of PBS alone was injected into the hock site of B6 mice (n=3). 30 min after injection, the mice were sacrificed, the hock and tibia tissue regions excised and flash frozen using liquid nitrogen. These samples were then submitted to the Southeast Center for Integrated Metabolomics at the University of for mass spectrometric analysis of kynurenine levels.

### Serum cytokine analysis

At the prescribed endpoints animals were placed under deep terminal anesthesia and cardiac punctures performed. 2 mL of blood were collected in microtainer serum separator tubes and centrifuged at 1500 rpm, for 5 minutes at room temperature. The serum was collected and store at −20°C. Cytokine analysis was performed via Luminex multiplex bead assay (Millipore) following manufacturer’s instructions.

### Imiquimod-induced Psoriasis

Pre-clinical modeling of psoriasis was carried out in 8-12-week-old female C57BL/6j (The Jackson Laboratory) mice in accordance with the Institutional Animal Care and Use Committee at the University of Florida. The model used was a modified version of that reported by van der Fits *et al*(*31*). Briefly, mice were anesthetized, and their backs were shaved, followed by application of depilatory cream to remove any remaining fur. Each day for 14 days total, 5% IMQ cream (62.5 mg, Patterson Veterinary Supply, cat. num. 07-893-7787) was applied to the backs of the mice. On the 3^rd^ day of IMQ application mice were subcutaneously injected with five 10 μg doses of IDO-Gal3 in sterile saline spread evenly throughout the back, or a sterile saline control (n = 12 per group). Disease severity was measured each day using a modified version of the Psoriasis Area and Severity Index (PASI) where area of effect is not taken into account. Erythema (redness), scaling, and thickening were scored independently and assigned a score on a scale of 0 to 4: 0, none, 1: slight, 2: moderate, 3: marked, 4: very marked. The cumulative score was reported as a measure of the severity of inflammation (scale 0-12).

### Near-infrared Conjugation and In Vivo Imaging of IDO-Gal3 Injected Subcutaneously

*In vivo* imaging of fluorescently tagged IDO-Gal3 was carried out in 8-12-week-old female C57BL/6j mice (The Jackson Laboratory) in accordance with the Institutional Animal Care and Use Committee at the University of Florida. Prior to injection, IDO-Gal3 was incubated with IRDye® 680RD NHS Ester (LI-COR, Inc., cat. num. 929-70050) according to the manufacturer’s instructions. Mice received a 40 μL (2.1 μg, 3.27 μM) subcutaneous injection to the hock of fluorescently labeled IDO-Gal3. Immediately following injection, and every subsequent 24 hours following injection, mice were imaged using a IVIS Spectrum In Vivo Imaging System (PerkinElmer). Fluorescent images were captured using the AF680 emission filter, subject size 1.5 cm, 0.2 s exposure time, field-of-view B (6.6 cm), medium binning (factor of 8) resolution, and a 1 F/Stop aperture. Relative fluorescent intensities were represented by a pseudo color scale ranging from red (least intense) to yellow (most intense).

### In Vivo Imaging of Tissue Distribution after Hock Injection

164 picomoles of NanoLuc-Gal3 in 40 μL of PBS were injected subcutaneously into the hock of B6 mice. At the prescribed time points, animals were euthanized in accordance with approved protocols. Organs and tissues of interest were harvested, weighed, processed, incubated with furimazine and bioluminescence quantified by a luminometer. Bioluminescence images were acquired using an IVIS Spectrum In Vivo System (IVIS). Living Image Software version 4.3.1 (Perkin Elmer, Waltham, MA) was used to acquire the data immediately after furimazine administration. Exposure time for the bioluminescence imaging was 1 second. Regions of interest were quantified as the average radiance (photons/second/cm2/sr). Specific amount of protein in tissue was determined by comparison with a standard curve of NanoLuc-Gal3 activity.

### Oral Listeria Infection after Hock Injection

Immune suppression from IDO-Gal3 was evaluated in response to oral infection with *Listeria monocytogenes* (strain EGDe). Tissue bacteria burden in the liver and spleen were used as a metric for infection where the liver is local for gut infection and the spleen represents systemic spread of infection. On the day prior to infection, 10-week-old naïve C57BL/6 mice (Taconic Biosciences) received IDO-Gal3 injection subcutaneously (hock), while control mice received sterile saline injection. The following day mice were orally infected with 2 x 10^9^ CFU/mouse of *Listeria monocytogenes*. To prepare for infection, bacteria were cultured overnight in BHI broth at 37 ^o^C, shaking at 220 RPM. Prior to infection a subculture of 2 mL of the bacteria and 18 mL BHI broth was cultured under the same conditions until reaching an OD_600_=0.8. Bacteria was pelleted, resuspended in 500 μL of sterile PBS, and 50 μL of bacteria was pipetted onto a small square of white bread and fed to each mouse individually. Once the entire piece of bread was eaten, mice were returned to their cage. At day seven post infection, mice were euthanized, followed by collection of the spleen, and liver. Tissue was homogenized, suspended in 1% saponin for one hour then plated at serial dilutions from undiluted to 1:1000. BHI Agar plates were treated with streptomycin to limit non-specific bacterial growth, as this strain of L*isteria monocytogenes* is streptomycin resistant. CFU’s were counted at 24 h post-plating.

### Murine Model of Periodontal Disease

All mice were lavaged with 25 μl of 0.12% chlorhexidine gluconate (3M) for three days. On day 4, 5, 6 and 7 mice received a 25 μl oral lavage with 2.5 × 10^9^ *Porphoromonas gingivalis* strain 381 and 2.5 × 10^9^. *Aggregatibacter actinomycetemocomitans* strain 29522 (ATCC) resuspended in 2% low viscosity carboxy-methyl-cellulose (Sigma-Aldrich). Oral lavage was repeated every week for 5 weeks. Each week, one day prior to the first day of infection (prophylactic) or one day after the last day of infection (therapeutic), 10ul of XXX IDO-Gal3 was injected into the submandibular space using a 30-gauge insulin syringe (Becton Dickenson). Each week of infection, on the first day of infection, prior to infection, microbial sampling of the oral environment was performed with calcium alginate swabs (Fisher Scientific). One week following the last infection, the mandibles were harvested to evaluate soft tissue soluble mediator expression, and bone morphometric analysis.

### Oral Soft Tissue Cytokine Expression

Mandibles with both soft tissue and bone were subjected to bead beating at two 2 min minute intervals with 2 min of cooling in between using 1.0 mm diameter zirconia silica beads (BioSpec) in cell extraction buffer (ThermoFisher) prepared with a protease inhibitor cocktail (mini cOmplete, Roche, Basel) and PMSF protease inhibitor (Abcam) to allow for dissociation and lysis of all soft tissue while leaving the hard tissues intact. MILLIPLEX® Multiplex Assays (EMD Millipore, Billerica, MA) were used to probe resulting lysates for IL6, IL1β, IL10 and MCP1 according to the manufacturer protocols. Data was acquired on a Luminex 200® system running xPONENT® 3.1 software (Luminex) and analyzed using a 5-paramater logistic spline-curve fitting method using Milliplex® Analyst V5.1 software (Vigene Tech). Data are presented as pg/ml normalized to total protein (BCA assay; ThermoScientific Pierce).

### Mandible Bone Morphometric Analysis

Mandibles were fixed in 4% buffered formalin for 24 h, stored in 70% alcohol and scanned at 18 μm resolution using a uCT system (Skyscan). 3D images were reconstructed, and the resulting images re-oriented spatially using anatomical landmarks with the NRecon and DataViewer softwares (Skyscan). A standardized 5.4 mm^3^ region of interest (ROI) was set with standardized dimensions of 1.5 (frontal) x 4.0 (sagittal) x 0.9 mm (transversal). Anatomical landmarks were used for the standardized positioning of the ROI: frontal plane = the roof of the furcation area between mesial and distal roots of the upper first molar; sagittal plane = anterior limit was the distal aspect of the mesial root of the first molar. The thickness of the ROI on the transversal plane was set to 50 slices (900 μm) and counted towards the palatal /medial direction beginning from the image that included the center of the upper first molar in its transversal width. A standardized threshold was set to distinguish between non-mineralized and mineralized tissues where total volume and total thickness of the ROI were calculated. *<u>Mean trabecular bone volume and thickness</u>*: The analysis assessed the percentage of mineralized tissue (BV; BT) within the total volume/thickness (TV; TT) of the ROI and is presented as presented the BV/TV ratio (mean trabecular bone volume) or BT/TT ration (mean trabecular thickness). *<u>Vertical bone loss</u>*: The distance from the cento-enamel junction (CEJ) to the alveolar bone was calculated at 12 sites over three molars and averaged to calculate the average vertical bone loss in mm.

### Oral Bacterial Burden

gDNA was isolated from microbial sampling of the oral environment using a DNeasy Kit (Qiagen) according to the manufacturers’ instructions. The gDNA was then probed for P. gingivalis 16S, A. actinomycetemocomitans 16S and total 16S using real time PCR. The percentage of A. actinomycetemocomitans 16S and P. gingivalis 16S within the total 16S compartment was calculated using the following formula: ct value of total 16S/ct value of A. actinomycetemocomitans or P. gingivalis 16S. 16s rRNA For: AGA GTT TGA TCC TGG CTC AG Rev: ACG GCT ACC TTG TTA CGA CTT; Pg For: CTT GAC TTC AGT GGC GGC AG; Rev: AGG GAA GAC GGT TTT CAC CA; Aa For: GTT TAG CCC TGG CCG AAG Rev: TGA CGG GCG GTG TGT ACA AGG.

### Mechanical Overloading Osteoarthritis Murine Model

IDO-Gal3 activity was assessed in a post-traumatic osteoarthritis mouse model adapted from Poulet *et al.* that applies cyclic mechanical loading to the knees of aged (6 months) mice, causing mechanical damage and consequent inflammation and cartilage degradation (*25, 26*). The mice were anaesthetized and placed in a fixture with the knee in flexion; loading (9 N) was applied axially for 500 cycles, and loading sessions are done on the mice 5 times per week during the experiment. IDO or IDO-Gal3 was prepared at 143 μM, and 20 μL of each treatment was injected intra-articularly at the start of each week of the study, with each knee receiving 4 total treatments. At the end of the 4 week study, mice were euthanized for analysis of gene expression and histopathology.

### Mechanical Overloading Osteoarthritis Pharmacokinetics Analysis

Pharmacokinetics of IDO and IDO-Gal3 retention after local injection at the disease site was assessed by intravital imaging over the course of 7 days. Both proteins were labeled with Li-Cor IRDye 680RD NHS ester (Li-Cor Biosciences, Lincoln, NE, USA) to visualize and measure protein knee retention. Mice were subjected to mechanical loading for two weeks before each treatment was administered via intraarticular injection. Intravital imaging was performed (Exc.: 672 nm, Emm.: 694 nm) immediately following injection, and every subsequent 24 hours. The fluorescence signal measured at the joint over time was normalized to the initial measurement for each knee, and an exponential decay was individually fit for each specimen. The area under the curve (AUC / bioavailability) for each joint was calculated from the best fit line of exponential decay. After 7 days, the mice were sacrificed and an ex vivo image was taken of each joint with the surrounding skin removed in order to increase measurement sensitivity.

### Mechanical Overloading Osteoarthritis Gene Expression Analysis

Gene expression was evaluated by TaqMan qPCR in the knee joint and the popliteal lymph node that drains the knee joint. Following sacrifice, knees and popliteal lymph nodes were excised. Combined joint tissue from the synovial wall and articular surface (not exceeding 30 mg total) and the popliteal lymph nodes were homogenized with bead pulverization in Qiazol. RNA was extracted and purified using the RNeasy Plus Mini Kit from Qiagen (Venlo, Netherlands) and quantified using NanoQuant plate from Tecan in a micro plate reader (Tecan Infinite 500, Tecan Group Ltd., Mannedorf, Switzerland). The RNA was coverted to cDNA using the iScript Synthesis Kit from Bio-Rad (Hercules, California, USA). Gene expression was calculated by the ΔΔCt method, normalizing to glyceraldehyde 3-phosphate dehydrogenase (GAPDH) and beta-actin (ACTB). TaqMan reagents were purchased from Thermofisher Scientific (Waltham, Massachusetts, USA) and used according to provided protocols, using appropriate primers (IL-12β: Mm01288989_m1, IL-6: Mm01210732_g1, MMP13: Mm00439491_m1, TNF-α: Mm00443258_m1, GAPDH: Mm99999915_g1, ACTB: Mm02619580_g1).

### Mechanical Overloading Osteoarthritis Histologic Staining and Scoring

Tissue samples were fixed in 10% formalin and decalcified in 20% EDTA for 7 days. A standard 8 h cycle of graded alcohols, xylenes and paraffin wax was used to process tissues before embedding and sectioning at 5μm thickness. Sections were mounted on positively charged glass slides and stained with H&E (hematoxylin and eosin) using the Gemini autostainer (ThermoFisher Scientific, Waltham, Massachusetts, USA). Safranin O staining was performed using the StatLab staining kit. Each joint was evaluated by at least two mid-frontal sections for both H&E and safranin O stains. A board-certified veterinary pathologist conducted histopathologic interpretations under blinded conditions.(*32*) OARSI scoring was based on medial and lateral tibial plateaus (scale of 0-6)(*33*), and a generic score was concurrently assigned based on H&E features and the safranin O staining of the tibial plateaus according to DJD methodology (Degenerative Joint Disease severity; scale 0-3).

### Surgically-Induced Osteoarthritis Rat Model

16 male Lewis rats (approximately 250 g) were acquired from Charles Rives Laboratories (Wilmington, MA, USA). Rats were acclaimed to the University of Florida housing facilities for 1 week. After acclamation, rats underwent baseline gait and von Frey behavioral testing. Following baseline behavioral testing, all rats received medial collateral ligament plus medial meniscus transection (MCLT+MMT) surgery to their right hind limb. Gait data was collected on weeks 3, 5, and 7 post-surgery, while von Frey testing was conducted weekly after surgery. At 8 weeks post-surgery, rats received a unilateral saline or IDO Gal-3 injection. Rats underwent von Frey testing the day following injection, and gait testing two days after injection. Both gait and von Frey data was collected weekly until euthanasia.

### Surgically-Induced Osteoarthritis Behavioral Study

MCLT+MMT surgery rats were anesthetized in a 2.5% isoflurane (Patterson Veterinary, Greeley, CO, USA) sleep box. Rats were then aseptically prepped with betadine surgical scrub (Purdue Products, Stamford, CT, USA) and 70% ethanol in triplicate, and transferred to a sterile field with anesthesia maintained via mask inhalation of 2.5% isoflurane. During MCLT+MMT surgery, a 1-2 cm midline skin incision was made along the medial aspects of the rat knee, and the skin was retracted to reveal the medial collateral ligament. The medial collateral ligament was transected and the knee was placed in the valgus orientation to stretch the medial compartment and expose the medial meniscus. The medial meniscus was cut radially, and absorbable 5-0 vicryl braided sutures (Ethicon, Somerville, NJ, USA) were used for muscle closure, and 4-0 ethilon nylon monofilament sutures (Ethicon, Somerville, NJ, USA) were used for skin closure. Rats recovered post-operatively in a warming box until weight bearing on all limbs. For pain management, rats received a subcutaneous injection of buprenorphine (0.05 mg/kg) (Patterson Veterinary, Greeley, CO, USA) intra-operatively and every 12 hours for 48 hours. Rats were grouped into a saline injection cohort (n=7) or IDO Gal-3 injection cohort (n=8). At 8 weeks after OA induction, rats received 30 µl unilateral injections of either sterile saline or IDO Gal-3 (33pg/µl) in the operated knee using sterile, allergy syringes (1 ml, 27G x 3/8) (Becton Dickinson and Company, Franklin Lakes, NJ, USA). First rats were anesthetized using 2.5% isoflurane (Patterson Veterinary, Greeley, CO, USA) and the operated knee was aseptically prepped as described above. Then the needle was inserted through the patellar ligament following the patellar groove into the joint space. The knee was flexed and the injection site was cleaned with sterile gauze and 70% ethanol.

### Surgically-Induced Osteoarthritis Tactile Sensitivity

Tactile sensitivity was assessed by measuring the 50% paw withdrawal threshold determined using the Chaplan up-down method for von Frey filaments^1^. Prior to surgery, animals were acclimated to the wire mesh-floored cage. Rats underwent von Frey testing the week before surgery, and weekly after surgery for 12 weeks. On week 8, rats were injected, then underwent von Frey testing the following day. During von Frey testing, a von Frey filament series (0.6, 1.4, 2.0, 4.0, 6.0, 8.0, 15.0, and 26.0 g) was applied to the plantar region of each hind foot. First the 4.0 g von Frey filament was applied. A less stiff filament was applied if a paw withdrawal occurred, and a stiffer filament was applied if a paw withdrawal did not occur. Using these data, the force at which rats were equally likely to withdrawal or tolerate was calculated via Chaplan’s approximation^1^.

### Surgically-Induced Osteoarthritis, Magnetic Capture

Commercially-available, streptavidin-functionalized particles (Life Technologies (Dynabeads MyOne™ Streptavidin C1, Cat. # 65001, Life Technologies, Carlsbad, CA, USA) were coated with biotinylated antibodies for CTXII (cat. # AC-08F1, ImmunoDiagnostic Systems, Copenhagen, Denmark), IL6, and MCP1 (cat. # 517703 and 505908, Biolegend, San Diego, CA, USA). Here, particles were washed 3 times in PBS, incubated for 2h on a tube revolver at room temperature in antibody mixes that contained either 8.9 ng/µL anti-CTXII, 8.9 ng/µL anti-IL6, or 357 ng/µL anti-MCP1, and then moved to static incubation at 4°C overnight. Particles were then washed three times in PBS containing 2% BSA and 2 mM EDTA (capture buffer), with final antibody amounts per particle measured to be 0.33 ng Ab/µg particle (anti-CTXII), 0.33 ng Ab/µg particle (anti-IL6), and 13.0 ng Ab/ µg particle (anti-MCP1).

Following euthanasia via exsanguination, magnetic capture was performed in both the operated and contralateral knee, as described in (*34*). Briefly, 300 µg of antibody-conjugated magnetic particles (equal parts for each particle type) were suspended in 10 µL of saline, then injected in the operated and contralateral knee. After 2 h incubation in the knee, particles were collected via 5 repeated 50 μl PBS washes of the knee, with collected fluid combined and particles in the fluid isolated via a magnetic separator. Collected particles were washed twice with capture buffer, then incubated for 15 min in the 100 mM Glycine-Tris buffer, pH 3.1, containing 2% BSA and 2 mM EDTA (release buffer). Following biomarker release, magnetic particles were isolated by magnetic separation, and the pH of the supernatant was adjusted to 8.3 for enzyme-linked immunosorbent assays (ELISA). CTXII was then measured in the supernatant using the Cartilaps CTX-II ELISA kit (cat. #AC-08F1, ImmunoDiagnostic Systems Cartilaps kit, Copenhagen, Denmark) according to the manufacturer’s instructions, and CCL2 was quantified using a rat CCL2 ELISA kit (Cat. # KRC1012 Life Technologies, Carlsbad, CA), according to the manufacturer’s instructions and modifications described in (*35*). IL-6 was quantified using ELISA developed in the laboratory using biotin anti-rat IL-6 antibody (cat. # 517703) and purified (coating) anti-rat IL-6 antibody (cat. # 517701, Biolegend, San Diego, CA, USA). Coating antibody was diluted in 100 mM NaHCO_3_, 34 mM Na_2_CO_3_ (pH 9.5), placed in coated microwells (Nunc MaxiSorp™, Cat. #434797, ThermoFisher Scientific Inc.), incubated for 30 min on a plate shaker then overnight at 4°C, washed 5 times with PBS with 0.5% Tween-20 (wash buffer), blocked for 1 h with PBS containing 2% BSA and 10% of heat-treated bovine serum, and finally washed 5 times with wash buffer. Samples and standards were pre-incubated with anti-IL-6 detection antibody (30 min at room temperature and then overnight at 4°C), then added to ELISA plate and incubated for 3 h. The plate was them washed 5 times, and 100 μl of avidin-HRP (cat. # 405103, Biolegend, San Diego, CA, USA, diluted 500 times in 2% BSA and 10% of heat-treated bovine serum) was added to the plate and incubated for 30 min. The plate was again washed 5 times, with 100 μl of tetramethylbenzidine (TMB) substrate added for 15 min, followed by 100 μl of stop solution (diluted sulfuric acid). Absorbance was read at 450 and 650 nm. Particles were quantified in 60 μl of capture buffer with particle suspensions read for absorbance at 450 nm using Synergy 2 Multi-Mode Microplate Reader, as described in (*34*).

### Surgically-Induced Osteoarthritis Histology

Following magnetic capture, operated and contralateral knees were dissected and placed in 10% neutral buffered formalin (Fisher Scientific, Pittsburgh, PA, USA) for 48 hours at room temperature. Following fixation, knees were decalcified using Cal-Ex (Fisher Scientific, Pittsburgh, PA, USA) for 5 days at room temperature, dehydrated through an ethanol ladder, and embedded in paraffin wax via vacuum infiltration. Then, 10 µm frontal sections were acquired, with at least one section taken at every 100 µm through the loading region of the medial meniscus. Slides were stained with toluidine blue.

### Surgically-Induced Osteoarthritis Joint Retention Study

16 male Lewis rats (approximately 250 g) were acquired from Charles River Laboratories (Wilmington, MA, USA). Rats were acclaimed to the University of Florida housing facilities for 1 week. Then rats underwent medial collateral ligament plus medial meniscus transection (MCLT+MMT) surgery. After 8 weeks, rats received a 50 µl injection of either Nano-Glo Luciferase (NL) or Nano-Glo Luciferase Galectin-3 (NL Gal-3). Rats were IVIS imaged immediately following injection, then 1, 2, 4, 8, 12, 16, 20, 24, and 28 days post-injection. After imaging on day 28, rats were euthanized, and both operated and contralateral knees were dissected for joint tissues distribution analysis via IVIS. In vivo joint retention was measured in the operated and contralateral knees of 16 male Lewis rats. At 8 weeks post MCLT+MMT surgery, knees were aseptically prepped and rats received bilateral injections of NL (50 µl, 3.27µM) (n=8) or bilateral injections of NL Gal-3 (50 µl, 3.27 µM) (n=8) in both operated and contralateral knees. (Surgeries and injections were conducted the same way as described above). Knees were flexed, then injected with 50 µl furimazine (50 µl, 1:50 dilution in PBS). Immediately after injection, knees were flexed again and luminescence was measured with IVIS using a 1 second (and 60 sec) exposure time in field of view D. Furimazine injections and IVIS imaging were repeated 1, 2, 4, 8, 12, 16, 20, 24, and 28 days post NL or NL Gal-3 injection. For analysis, a region of interest (ROI) was drawn around the largest luminescent signal and copied to create an identical sized ROI for all knees.

After imaging on day 28, rats were euthanized via cardiac puncture under deep isoflurane anesthesia. Knees were dissected to isolate the patellar tissue, tibial tissue, femoral tissue, and meniscus. The patellar tissue included the patella, patellar synovium, and fat pad. The tibial tissue included the tibia and attached synovium. The femoral tissue included the femur and attached synovium. Tissues were placed in 24 well plates and incubated with furimazine (1:50 dilution in PBS) at room temperate for roughly 1 minute. After incubation, tissues were removed from the 24 well plates and luminescence was measured with IVIS using auto-exposure in field of view C. For assessment, individual ROIs were drawn around the patellar tissue, tibial tissue, femoral tissue, and meniscus. The ROI for each tissue was copied to create an identical sized ROI for the respective tissue.

### Statistical Analysis

Statistical analyses were performed using Prism Software (GraphPad Software, Inc.; La Jolla, CA), with the following exceptions. Surgically-induced osteoarthritis data was analyzed using RStudio, and MATLAB was used for Mann Whitney U-Test on psoriasis data. Study-specific analyses are reported in figure captions.

## Supplementary Information for

This Supplementary Information document includes:

**Supplementary Figure 1:**
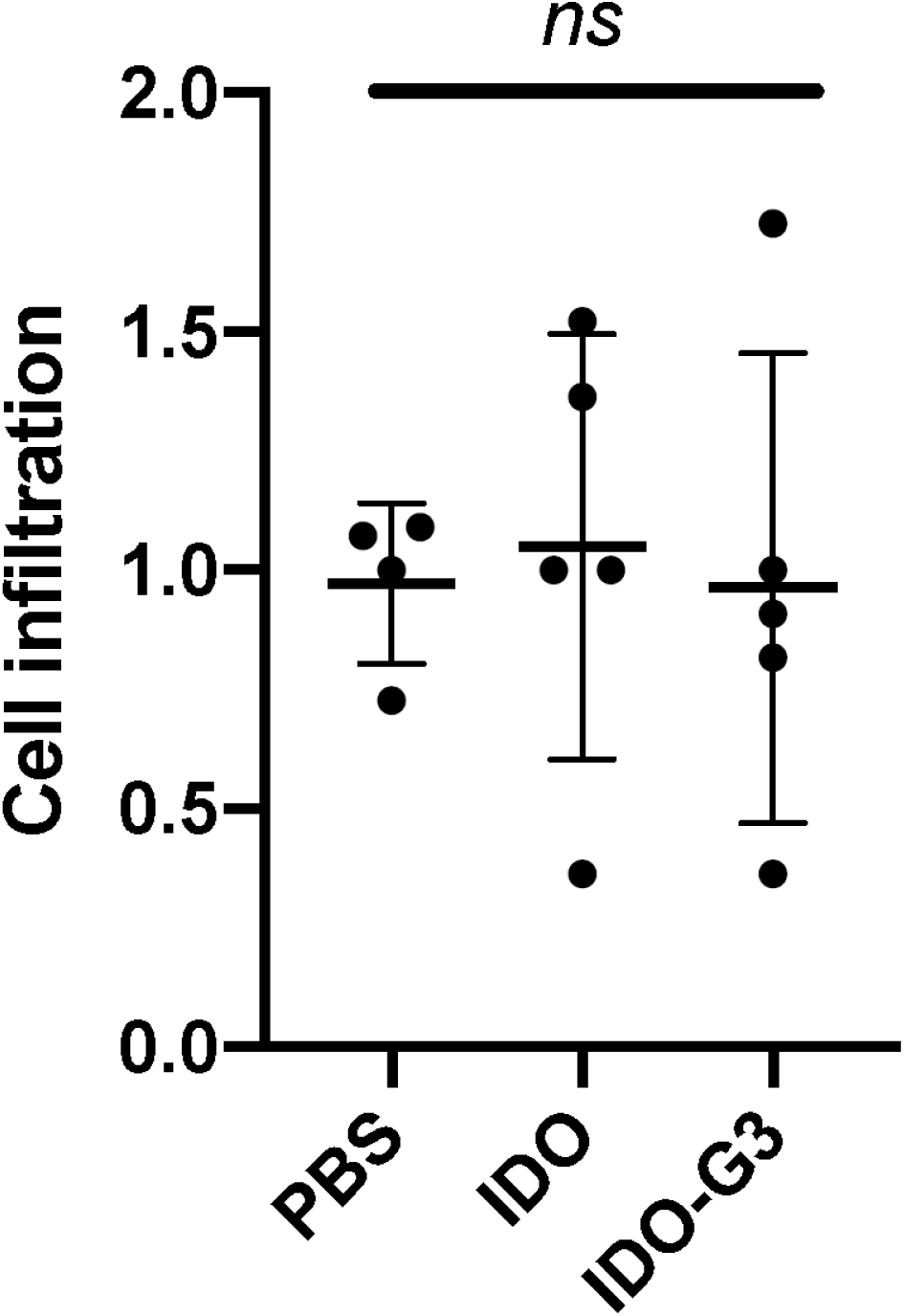
Quantitative analysis of H&E histology images showing that subcutaneous injection of IDO or IDO-G3 alone do not increase cell infiltration compared to vehicle injection alone. “ns” indicates p > 0.05, ANOVA with Tukey’s post-hoc analysis.

**Supplementary Figure 2:**
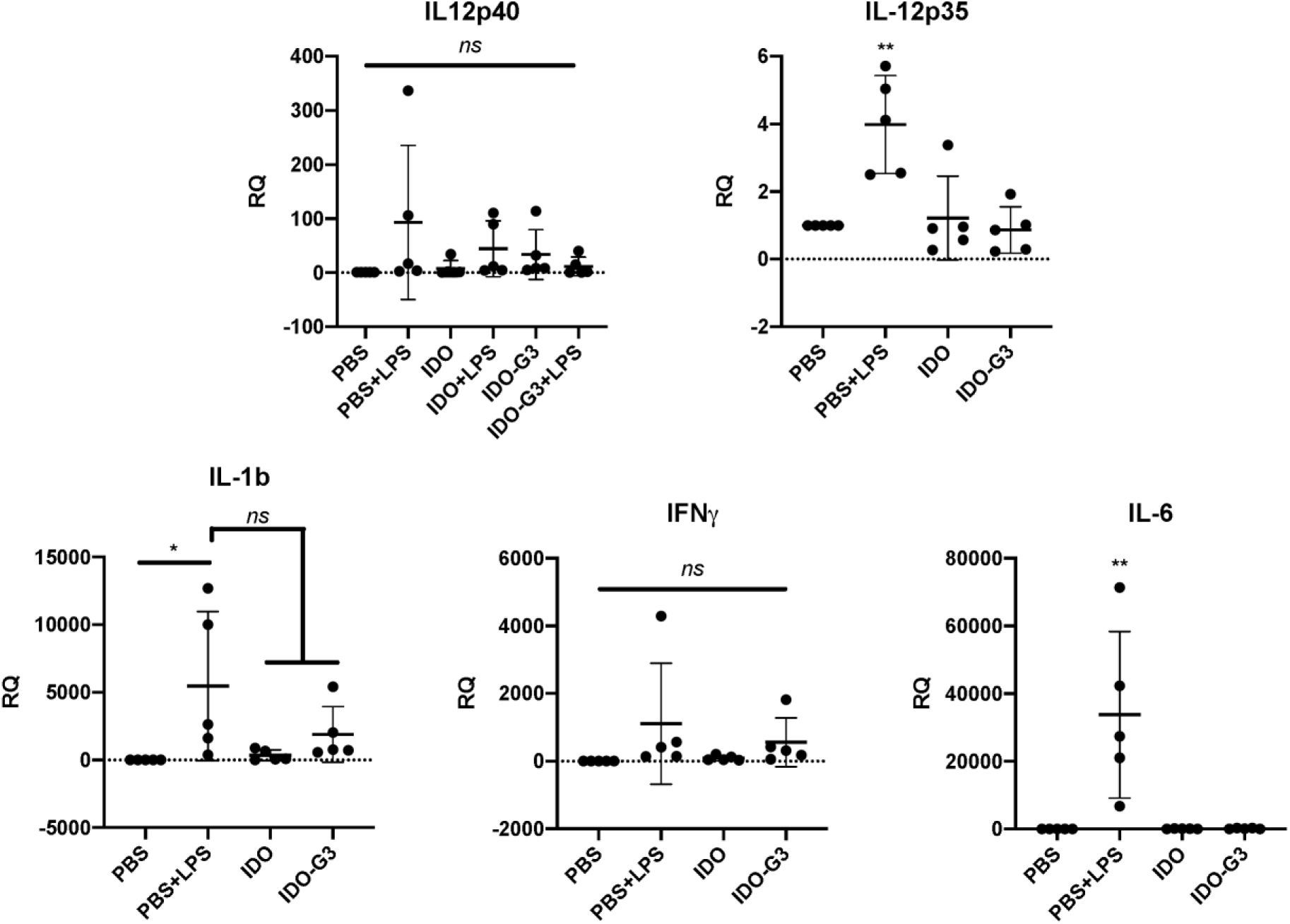
Cytokine gene expression following subcutaneous injection of vehicle (PBS), IDO, or IDO-G3 at t = 0 and then vehicle or LPS (labeled as “+LPS”) at t = 24. * represents p < 0.05, ** represents p < 0.01 compared to all other groups, unless otherwise indicated, ANOVA with Tukey’s post-hoc.

**Supplementary Figure 3.**
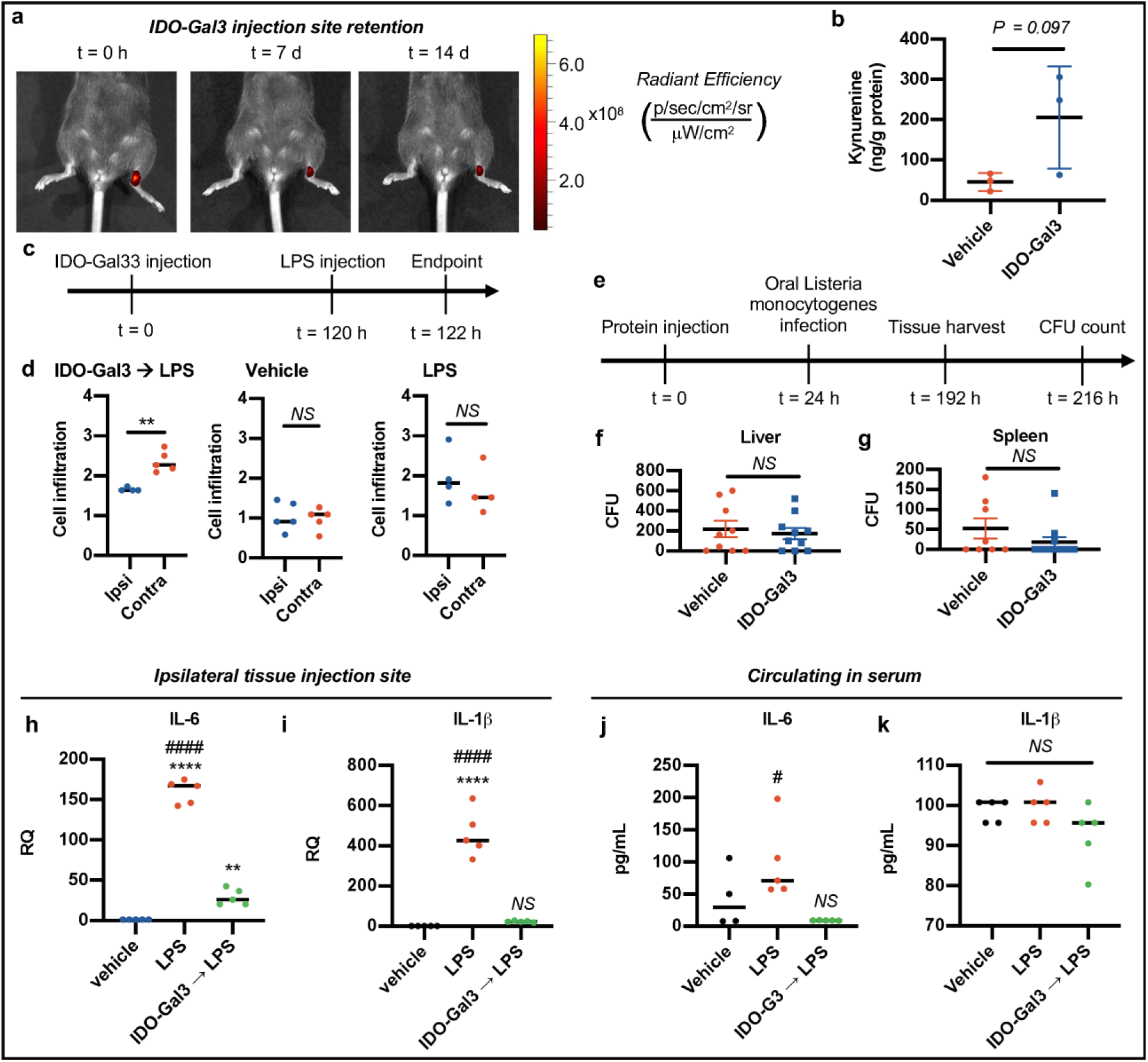
IDO-Gal3 functions locally and is not systemically suppressive. (a) In vivo imaging of fluorophore-labeled IDO-Gal3 injected subcutaneously (s.c.) in the hock demonstrating tissue retention out to 14 d. (b) Mass spectrometry analysis quantification of kynurenine levels in hock and surrounding tissue indicating in vivo IDO enzymatic activity. (c-d) LPS challenge 120 h after s.c. hock injection at both ipsilateral (IDO-Gal3 pre-treated) and contralateral (vehicle) sites yields a local ipsilateral reduction in inflammatory cell infiltration while maintaining a robust contralateral inflammatory response. (e-g) Administering IDO-Gal3 into the s.c. hock did not alter clearance of Listeria monocytogenes after oral infection challenge, in the liver or spleen compared to vehicle control. Hock s.c. IDO-Gal3 pretreatment 120 h (c) suppressed transcript levels of IL-6 (h) and IL-1β (i) at the ipsilateral LPS challenge site, blocking systemic increase in IL-6 protein (j) and maintaining baseline systemic IL-1β (k), quantified in serum by mass spectrometry. Statistical analyses: (b) Student’s t-test, n = 3; (d) Student’s t-test, two symbols denotes p < 0.01, n = 5;, (f) Student’s t-test, “NS” indicates no difference, two symbols denotes p < 0.01; (h-k) One-way ANOVA with Tukey’s post-hoc. “NS” indicates no difference, one symbol denotes p < 0.05, four symbols denotes p < 0.0001, n = 5.

**Supplementary Figure 4:**
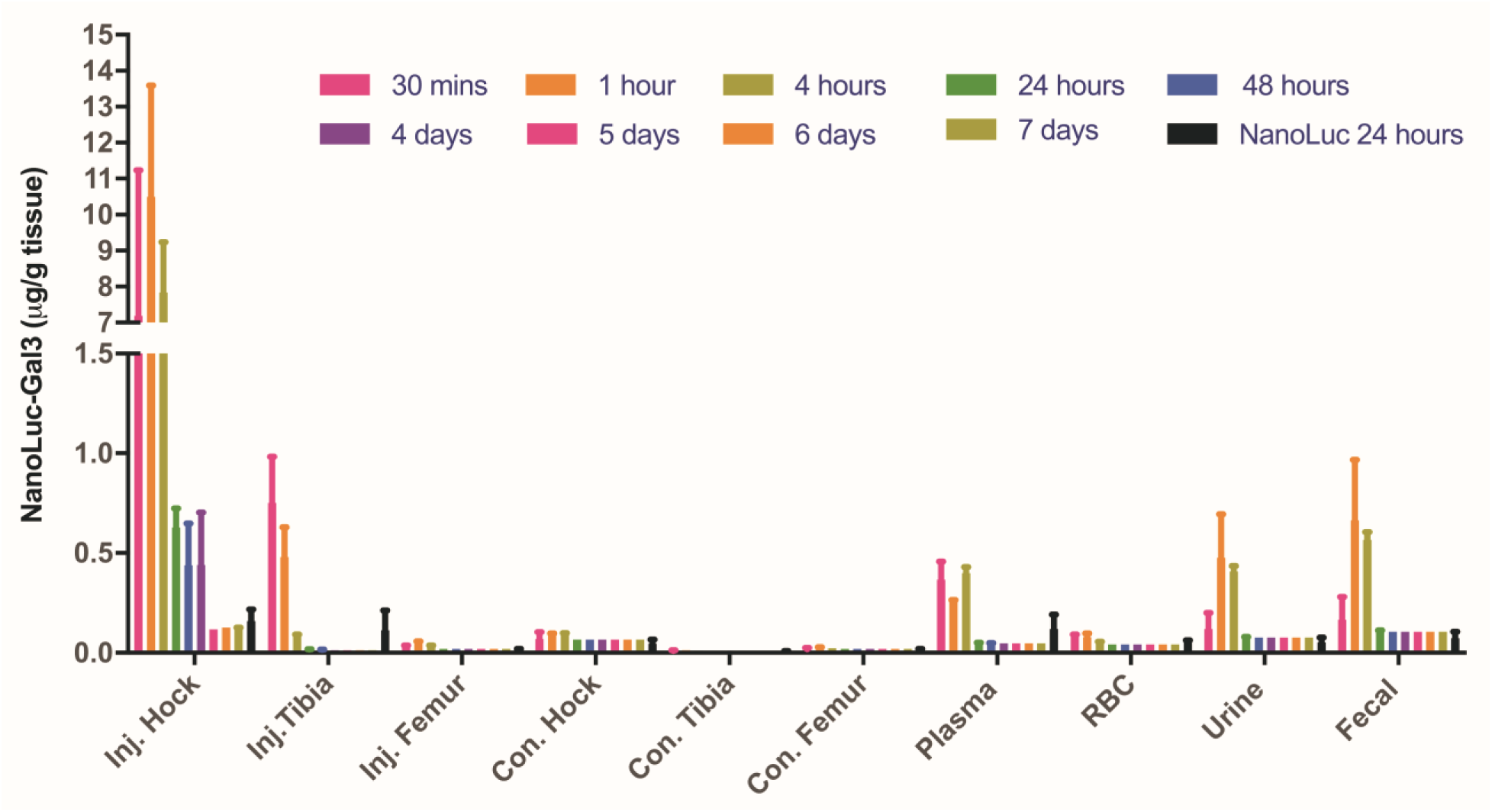
Biodistribution of NL-G3 determined from NL bioluminescence at various time points following subcutaneous injection into the hock at t = 0.

**Supplementary Figure 5:**
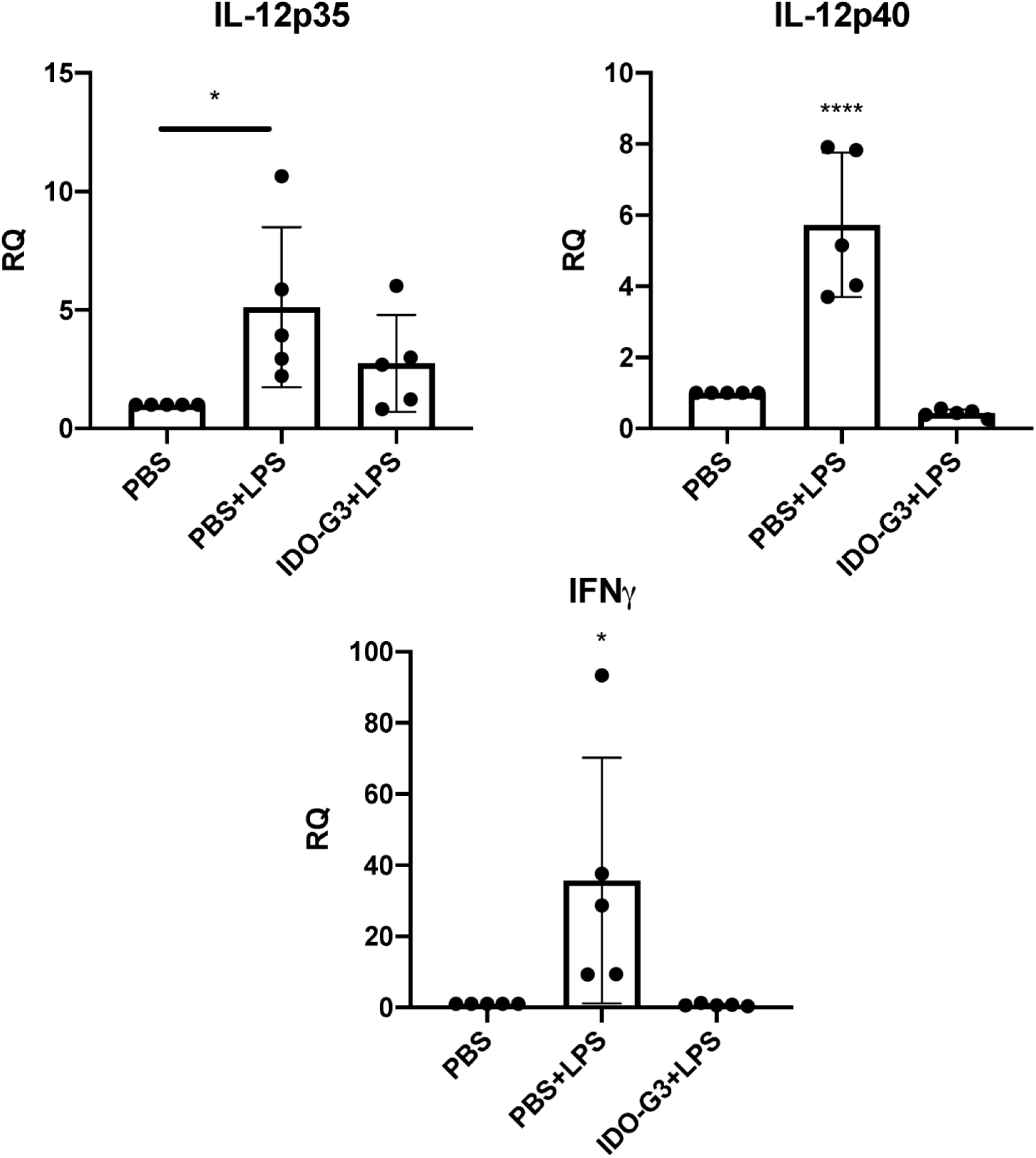
Cytokine gene expression after subcutaneous injection of vehicle (PBS) or IDO-G3 at t = 0 followed by ipsilateral injection of vehicle or LPS at t = 120 h. * represents p < 0.05, **** represents p < 0.0001 between all groups relative to PBS + LPS, unless otherwise specified, ANOVA with Tukey’s post-hoc.

**Supplementary Figure 6:**
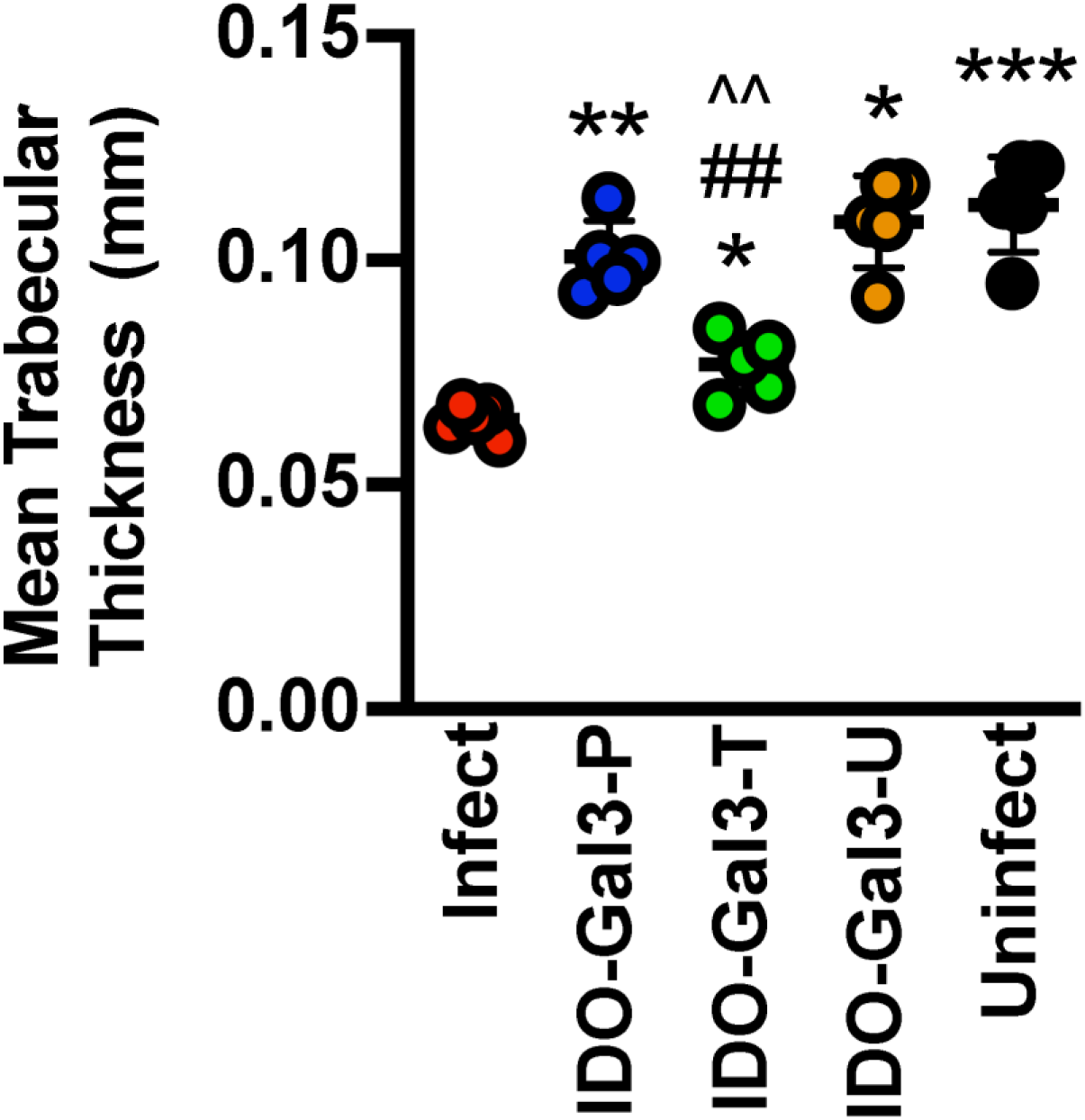
Mean trabecular bone thickness calculated from microCT images following prophylactic (IDO-Gal3-P) or therapeutic (IDO-Gal3-T) submandibular injection of IDO-G3 in mice infected with PG and AA. “IDO-Gal3-U” denotes animals that received IDO-G3 but were not infected, “Infect” denotes untreated infected mice, and “Uninfect” denotes uninfected and untreated mice. * indicates statistically significant differences compared to “Infect”, # indicates statistically significant differences compared to “Uninfect”, and ^ denotes statistically significant differences of IDO-Gal3-P relative to IDO-Gal3-T. One symbol denotes p < 0.05, two symbols denotes p < 0.01, three symbols denotes p < 0.001, and four symbols denotes p < 0.0001, ANOVA with Tukey’s post-hoc.

**Supplementary Figure 7:**
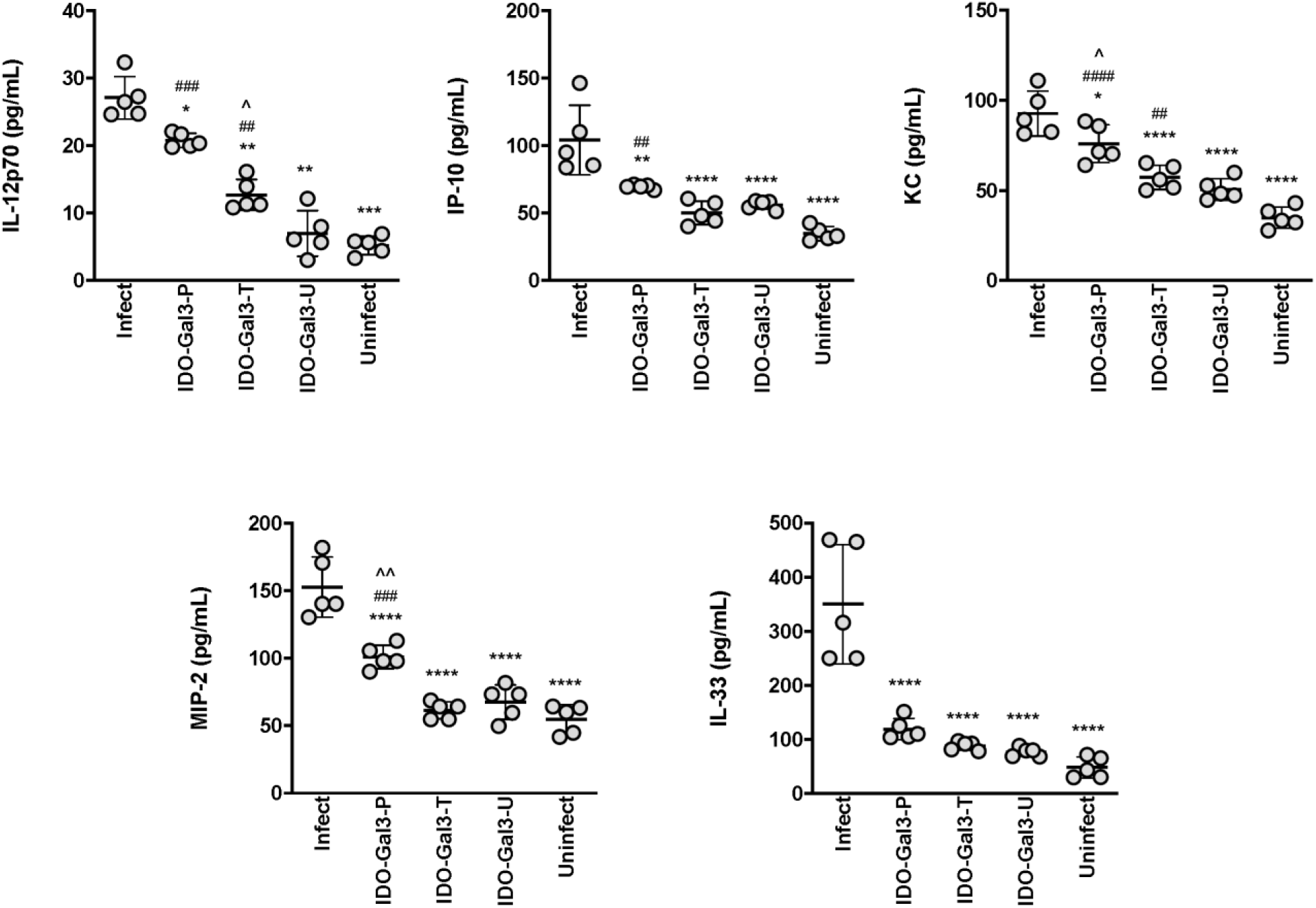
Local cytokine expression following prophylactic (IDO-Gal3-P) or therapeutic (IDO-Gal3-T) submandibular injection of IDO-G3 in mice infected with PG and AA. Both prophylactic and therapeutic administration of IDO-Gal3 reduced gingival inflammatory protein levels of IL12p70, IP10, KC, MIP2 and IL33 with more pronounced effect following therapeutic administration. “IDO-Gal3-U” denotes animals that received IDO-G3 but were not infected, “Infect” denotes untreated infected mice, and “Uninfect” denotes uninfected and untreated mice. * indicates statistically significant differences compared to “Infect”, # indicates statistically significant differences compared to “Uninfect”, and ^ denotes statistically significant differences of IDO-Gal3-P relative to IDO-Gal3-T. One symbol denotes p < 0.05, two symbols denotes p < 0.01, three symbols denotes p < 0.001, and four symbols denotes p < 0.0001, ANOVA with Tukey’s post-hoc.

**Supplementary Figure 8:**
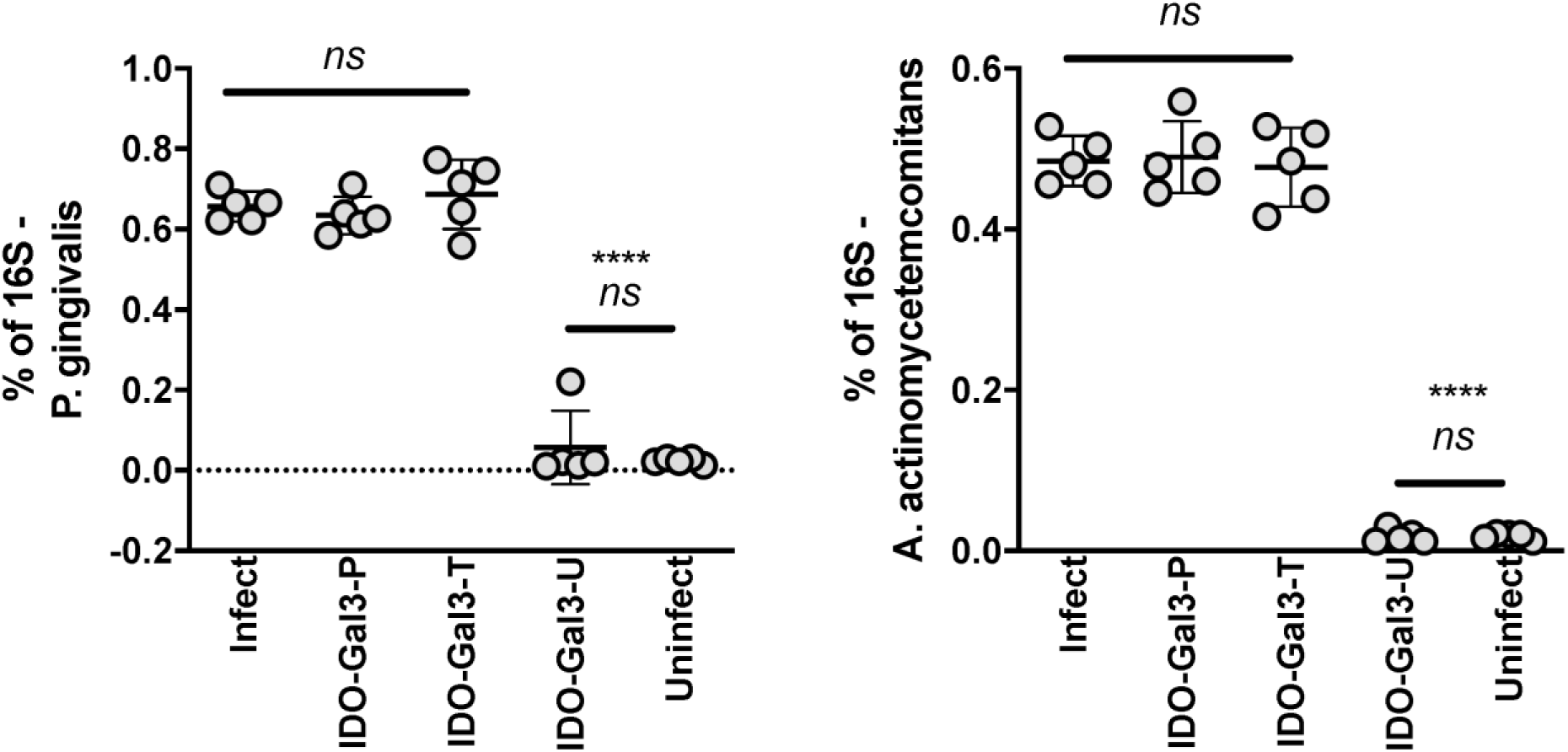
Bacterial load in infected mice that received a prophylactic (IDO-Gal3-P) or therapeutic (IDO-Gal3-T) submandibular injection of IDO-G3. Recovery of *P. gingivalis* and *A. actinomycetemcomitans* from the oral cavity as measured by qPCR using 18s specific primers. Data are expressed as a % of total bacteria recovered as measured by qPCR using 18s consensus primers. Neither prophylactic (IDO-Gal3-P) nor therapeutic (IDO-Gal3-T) administration of IDO-Gal3 affects the % of *P. gingivalis* and *A. actinomycetemcomitans* recovered from the oral cavity. “IDO-Gal3-U” denotes animals that received IDO-G3 but were not infected, “Infect” denotes untreated infected mice, and “Uninfect” denotes uninfected and untreated mice. “ns” indicates no difference between indicated groups, **** indicates p < 0.0001 compared to “infect” group, ANOVA with Tukey’s post-hoc.

**Supplementary Figure 9:**
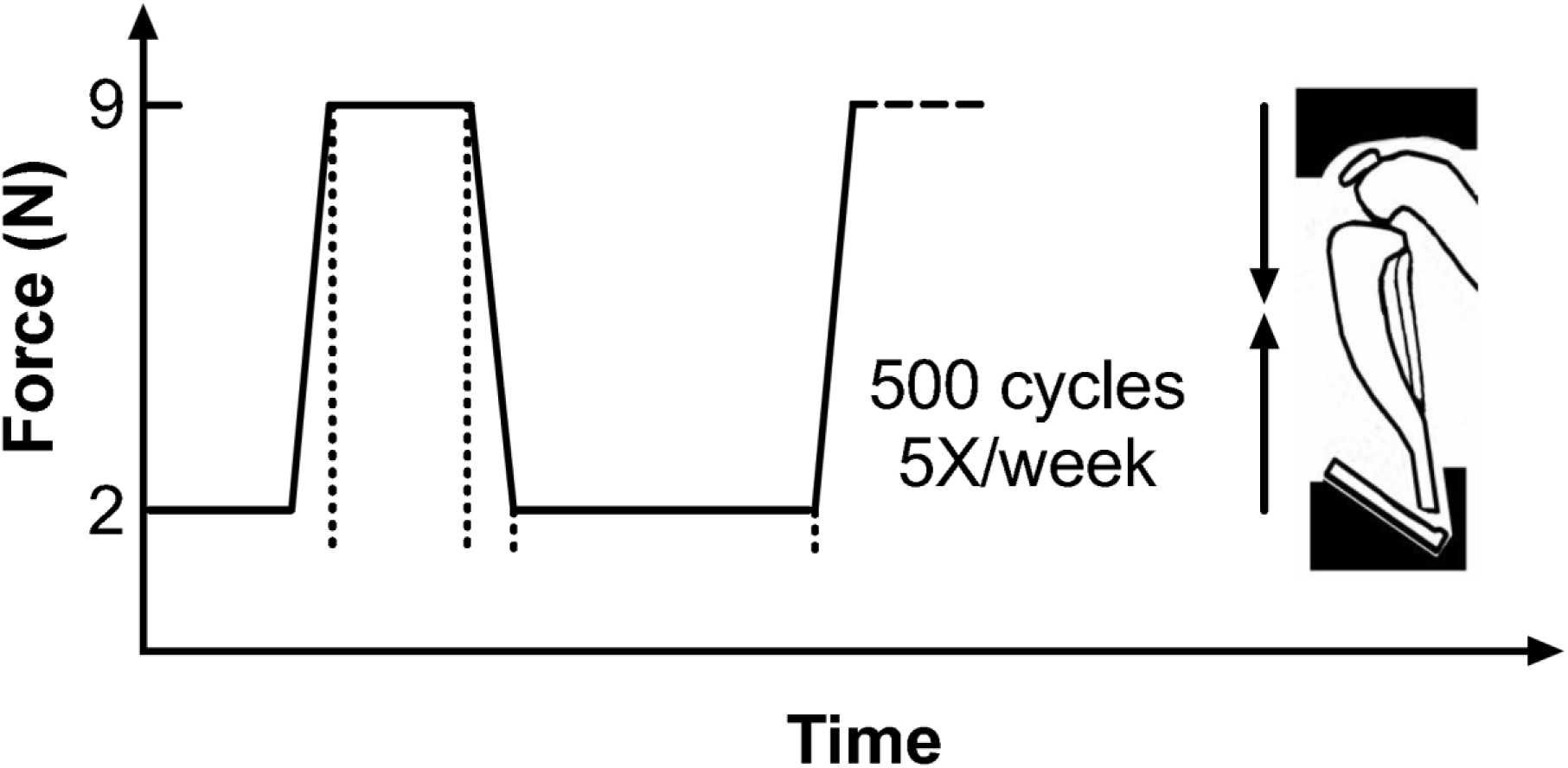
Graphical representation of the mechanical over-loading protocol used to induce murine osteoarthritis.

**Supplementary Figure 10:**
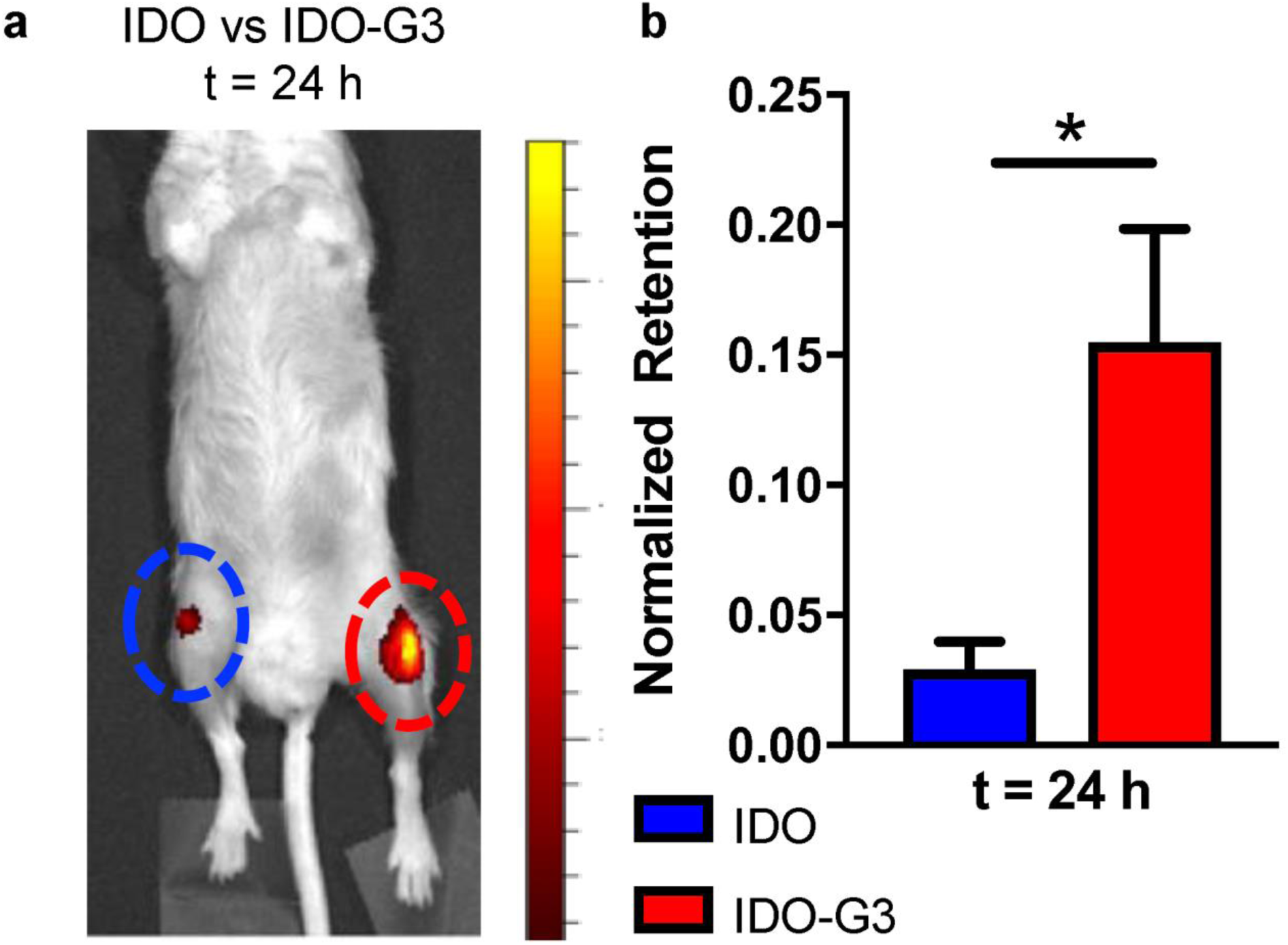
(a) Representative image of IDO (blue, left) and IDO-G3 (red, right) retained within the mouse knee 24 h after injection. (b) Fraction of initial fluorescence signal retained at the injection site 24 h after injection. * represents p < 0.05, ANOVA with Tukey’s post-hoc.

**Supplementary Figure 11:**
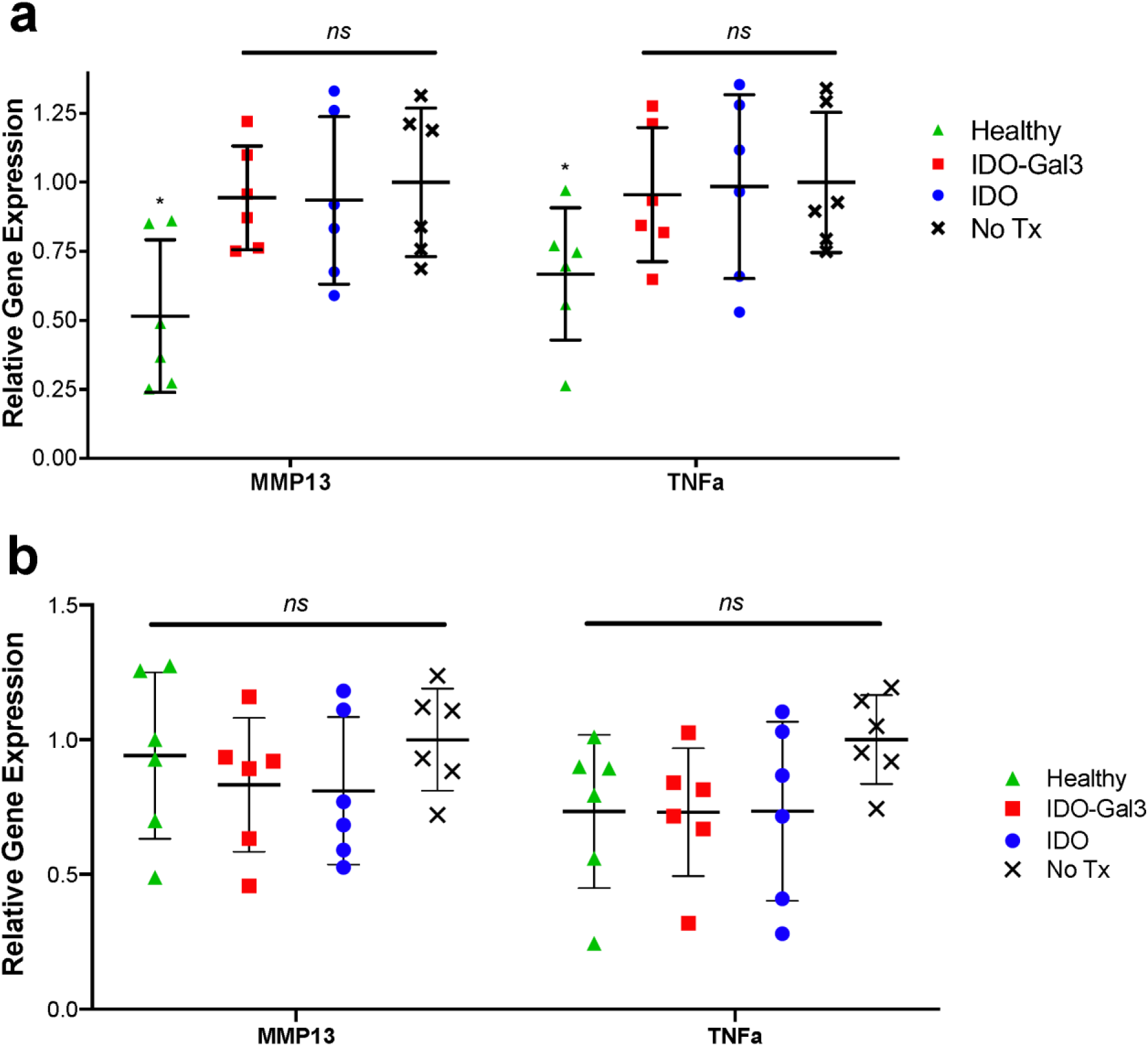
Gene expression in (a) the knee with mechanically-induced osteoarthritis and (b) the popliteal lymph node. * denotes p < 0.05 relative to all other groups, “ns” denotes no difference between indicated groups, ANOVA with Tukey’s post-hoc.

**Supplementary Figure 12:**
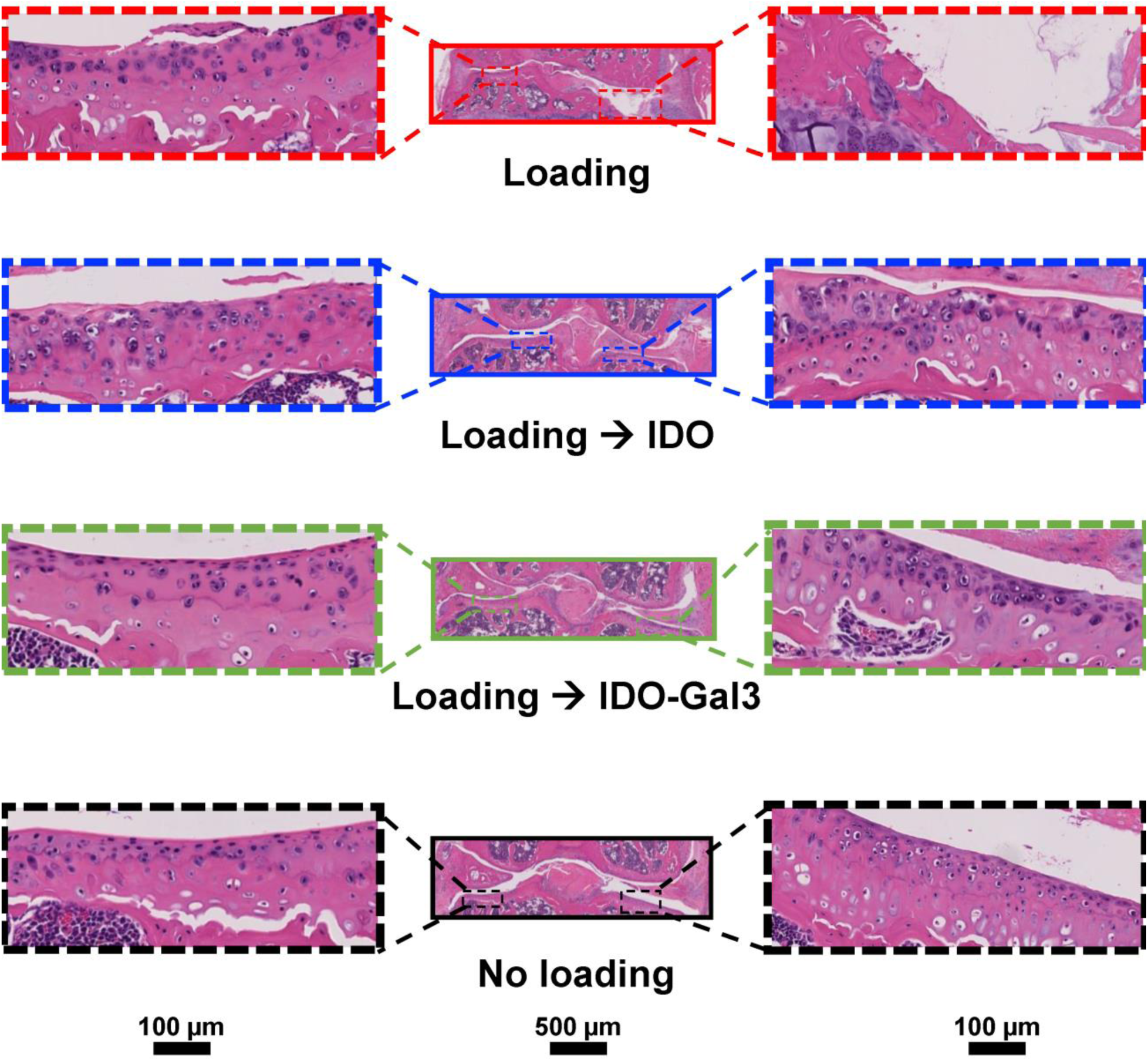
Representative Hematoxylin & Eosin staining of whole joints and tibial articular cartilage surfaces per treatment group shown as a serial magnification.

**Supplementary Figure 13:**
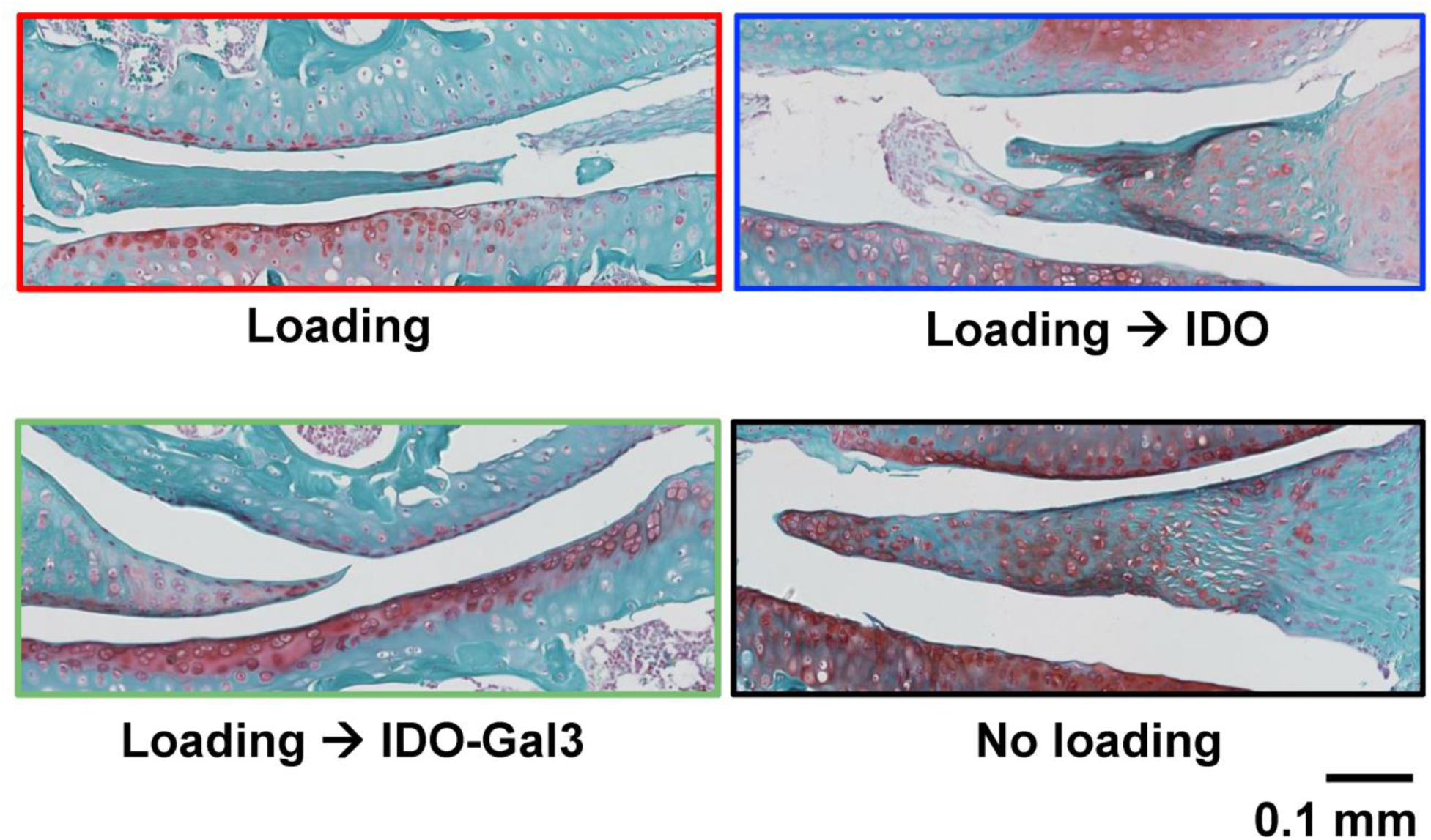
Representative Safranin-O/Fast Green staining of the femoral-tibial cartilage interface with accompanying meniscus per treatment group.

**Supplementary Figure 14:**
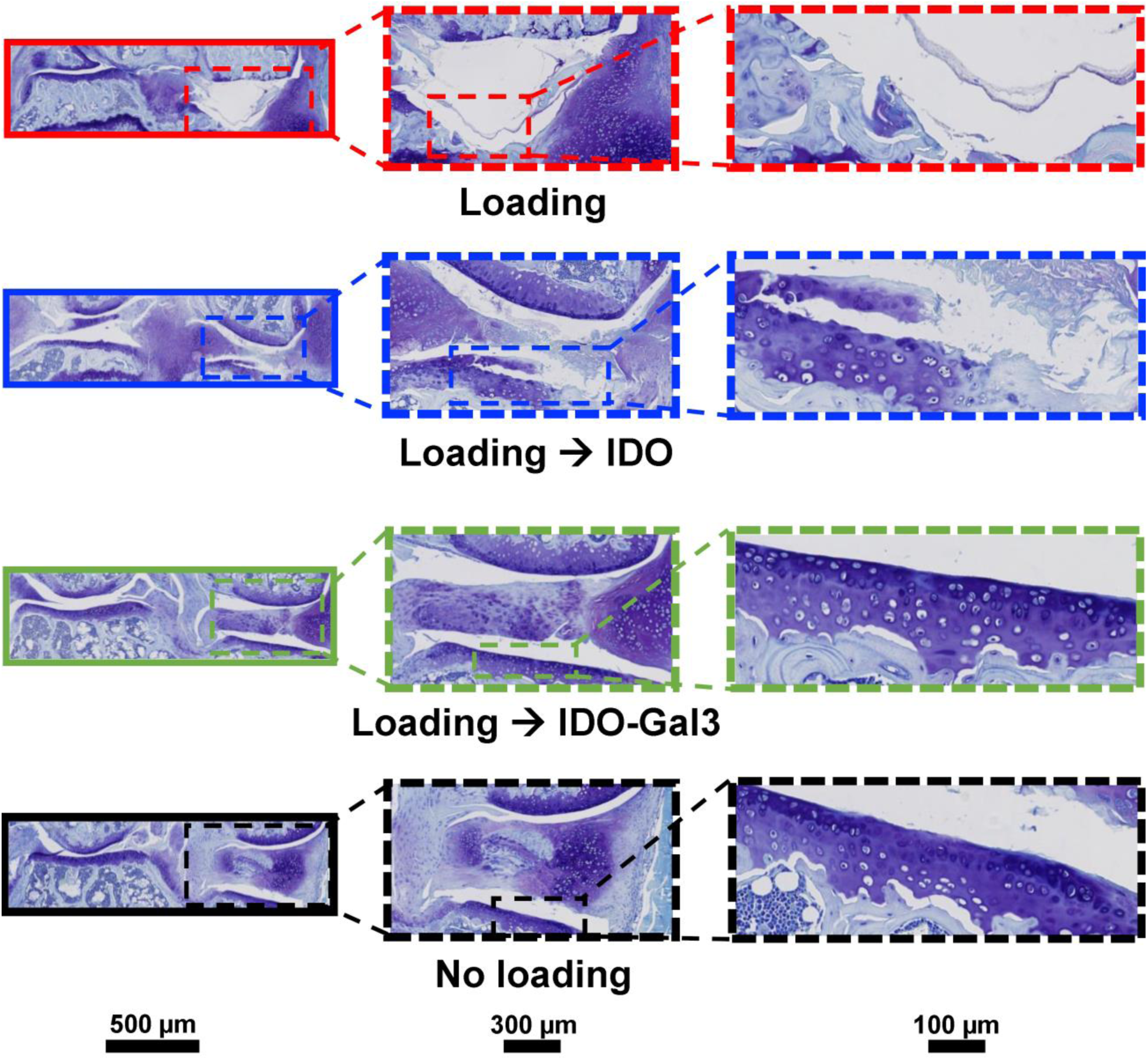
Representative Toluidine Blue staining of whole joints, half joints, and tibial articular cartilage surfaces per treatment group shown as a serial magnification.

**Supplementary Figure 15:**
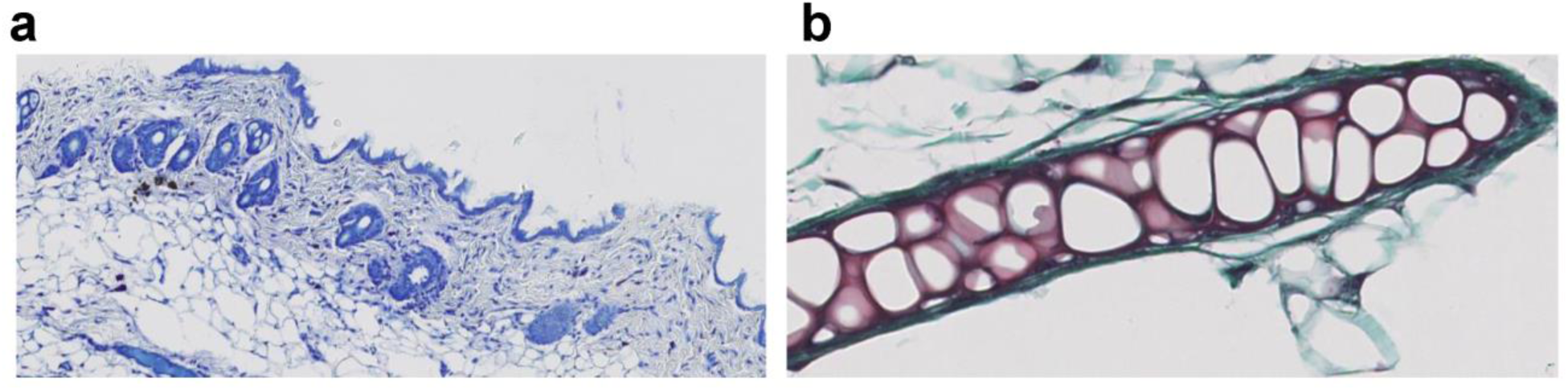
(a) Toluidine staining control of mouse skin containing mast cells. (b) Safranin-O/Fast green staining control of mouse ear tissue.

**Supplementary Figure 16:**
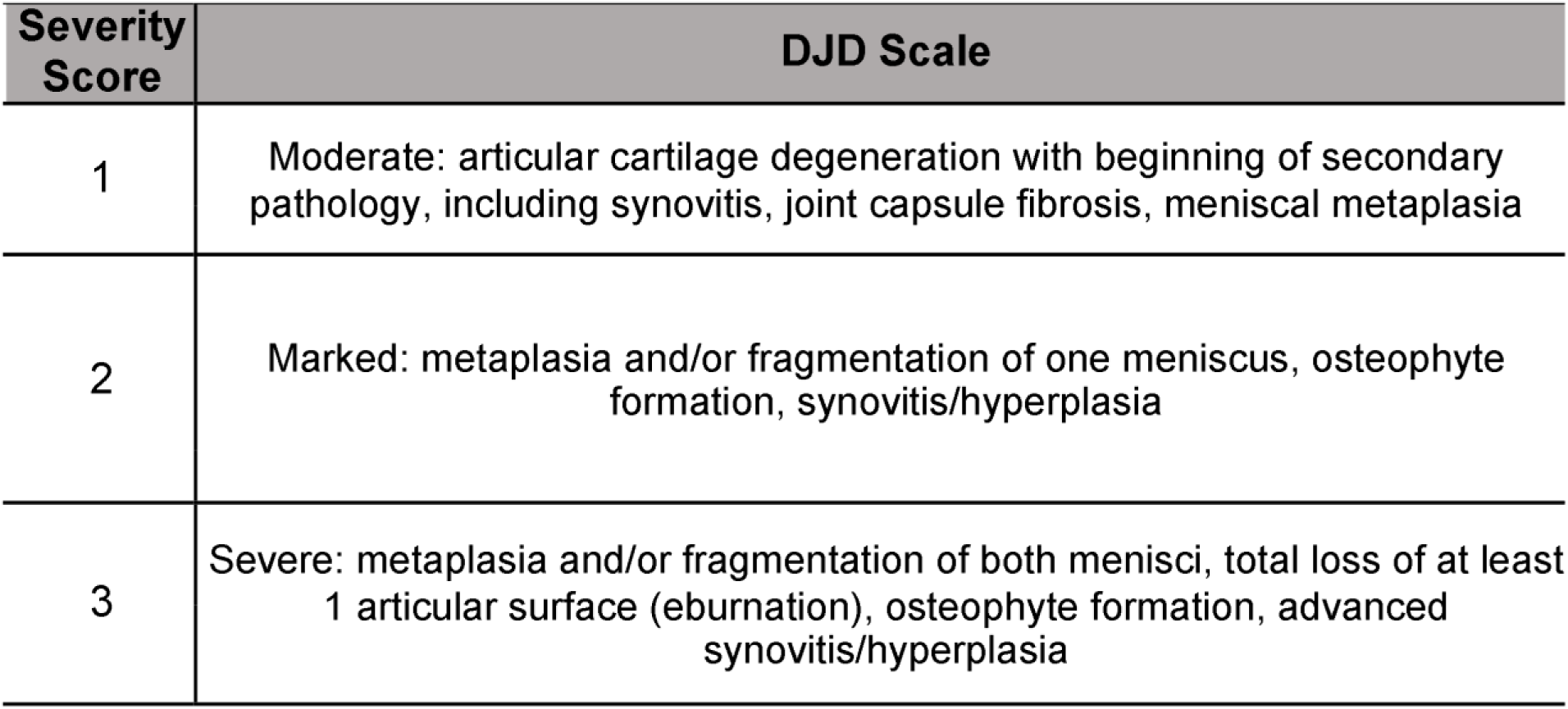
Overview of the DJD scoring rubric.

**Supplementary Figure 17:**
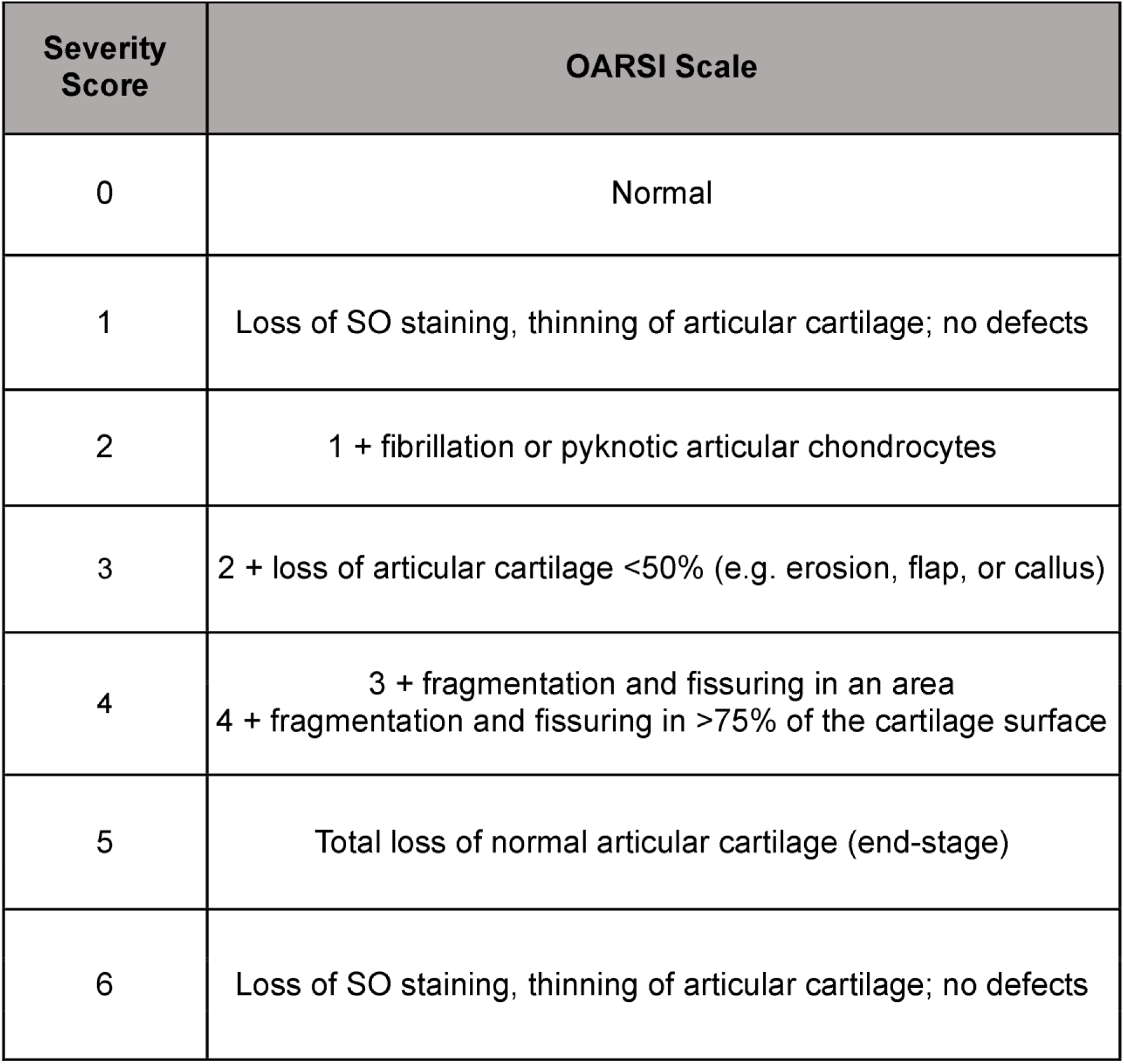
Overview of the OARSI scoring rubric.

**Supplementary Figure 18.**
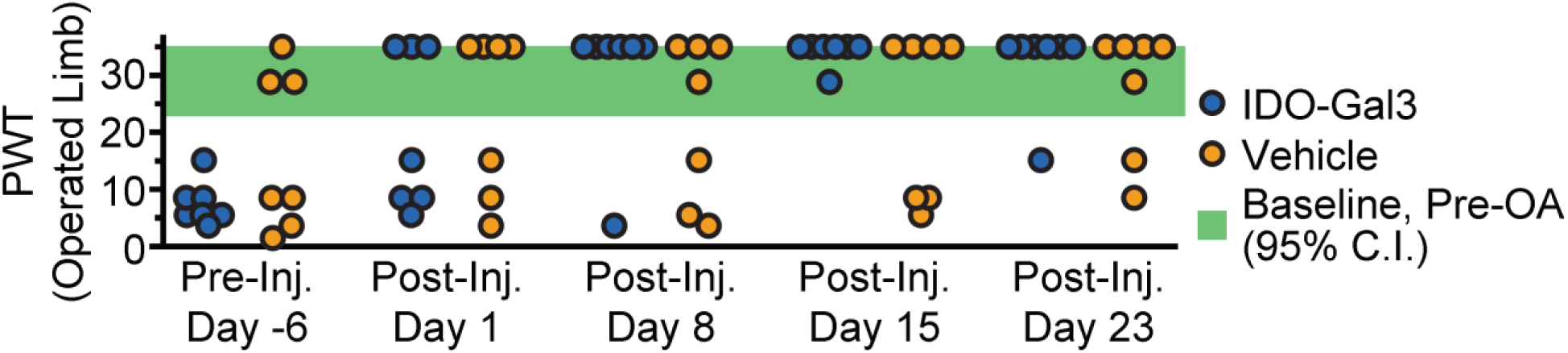
Paw withdrawal thresholds (PWT) where the animal is equally likely to tolerate versus withdraw from on touch stimulus.

**Supplementary Figure 19.**
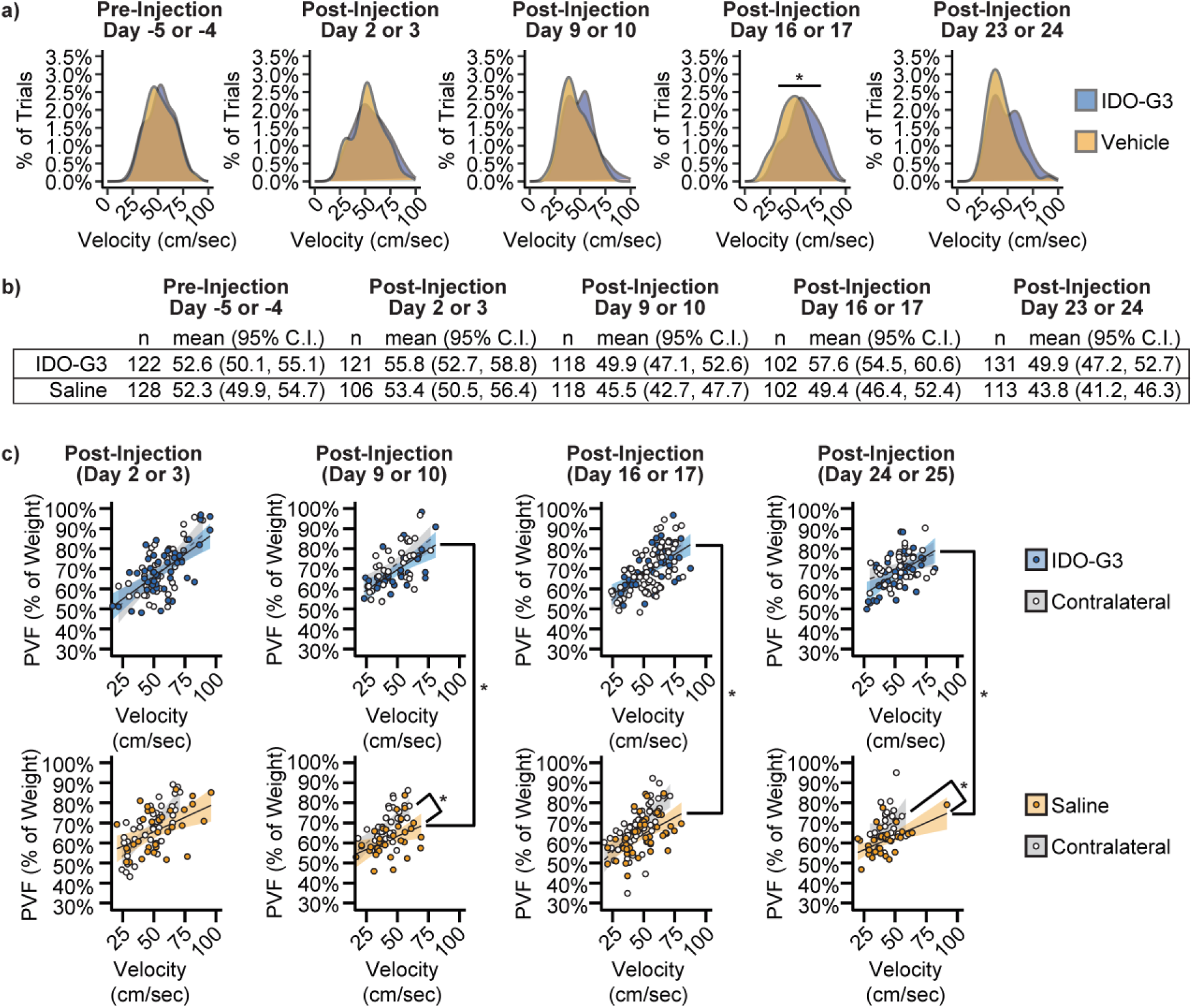
Distribution of selected walking velocity (a) along with trial counts, mean, and 95% confidence intervals (b). IDO-Gal3-treated animals used faster walking velocities at post-injection day 16 (p=0.02) and tended to use faster velocities at post-injection day 23 (p=0.051). Raw data of the peak vertical force-time relationships used to project 95% confidence intervals for IDO-Gal3- and saline-treated knees, along with their respective contralateral controls.

